# Examining gender imbalance in chemistry authorship

**DOI:** 10.1101/2020.07.08.194035

**Authors:** Adam D. Cotton, Ian B. Seiple

## Abstract

The gender gap in chemistry has been the topic of much debate, with many perspectives stating that the field has improved, and outdated sexist views are behind us. While these views are common, we wanted to assess the accuracy of these comments from a data-driven perspective. In this study, we use PubMed to obtain the names of first and last authors for every paper published since 2005 across 15 journals. Each name was cross-referenced with a name-based gender API to give a predicted binary gender and a confidence score based on population data. We show that historically there has been an extensive overrepresentation of men in both first and corresponding authorship, and that there is no strong trend towards parity since 2005. We demonstrate that papers with female corresponding authors have more equitable gender representation of first authors. Finally, we find that there is significant variability among journals in the gender make-up of their editorial boards. We hope this analysis spurs creative discussions on how we can improve equitable gender representation in chemistry publications.

## Introduction

In recent weeks there has been extensive conversation regarding gender diversity in chemistry, spawned in part by an article published online (and then removed and erased from the website) in *Angewandte Chemie* entitled “‘Organic synthesis–Where now?’ is 30 years old. A reflection on the current state of affairs”. Commentaries since its publication have stated our field has improved and outdated sexist views are behind us, we wanted to assess how accurate these perspectives are using a data-driven approach. Our goal is to provide data to the community to assist in constructive conversations about how we can progress. There have been numerous validated studies demonstrating how diversity in teams improves creativity and productivity^1,2^. Any deviation of gender distribution in a given field from the abundance in the general population is caused by systemic biases.

We wrote a python-based web scraper to extract the first (given) name of the first and last (corresponding) authors from papers published on PubMed each year since 2005. 2005 was chosen due to the lack of quality data in prior years. We focused our efforts on organic chemistry journals, the closely related chemical biology field, and Nature, Cell, Science (NCS). In addition to authors’ names, we obtained the country of the corresponding author and assumed this would be the same for the first author. Self-identification is the only accurate way to assign gender, but such data is not currently widely available for authors in chemistry journals. In the absence of sufficient self-identification data, and given that many first names strongly correlate with gender, we sought to use first names to predict gender ratios probabilistically. We utilized <u>Gender API</u>^5^ to determine a gender probability and confidence score for each name, using country to improve accuracy for common names. Due to the binary nature of the API used, we could only account for two genders: male and female. There were occasions where the API couldn’t interpret the name or where the name was not in the database, and these instances were removed from the analysis. Gender API outputs an accuracy score and sampling size, we set strict data quality requirements that were used to calculate population parameters. Any names with <95% accuracy score were excluded to minimize uncertainty. Any years for a journal with a high-quality data set of less than 25 were removed from any further analyses. Of 392,754 authors, we removed 139,505 that did not meet the inclusion criteria. Details, graphical figures, individual journal graphs, and further discussion about the limitations of our approach can be found in the supporting information. All Python scripts used are available on <u>github</u>^6^.

## Results

The first question we assessed was if there has been a gender disparity for first authors and for corresponding authors, cumulatively over the past 16 years, and if there were any dissonance between different journals. It quickly became clear that there has historically been exceedingly low representation of women in Chemistry (Figure 1). In *Journal of the American Chemical Society* (JACS), a leading chemistry journal, women have represented 9.5% of corresponding authors and 21.4% of first authors (Figure 1). These values were consistent across the different chemistry journals. Interestingly medicinal chemistry journals such as *Journal of Medicinal Chemistry* (J Med Chem) had a higher representation of women, who made up 15.7% of corresponding authors and 29.9% of first authors. We speculate that these values may arise due to the many different contributing factors to authorship in academia versus industry. Due to this difference, medicinal chemistry journals were separated from the other traditional chemistry journals. Chemical biology is a closely related field to Chemistry, but gender ratios in the three chemical biology journals included in this study grouped separately from chemistry journals. ACS Chemical Biology had the highest representation of women, who comprised 18.4% of corresponding authors and 34.6% of first authors. It is important to note that these values are far from representative of the gender demographics seen in the general population. Interestingly the NCS journals separated in two groups throughout our analyses, with Nature and Science closely correlating with each other while Cell demonstrated female to male ratios that were closer to parity. Finally, it is apparent there is a large disparity between first authors and corresponding authors, which tracks with faculty and student gender percentages; 39% of chemistry graduate students and only 12% of faculty are women. This aligns with the documented decline of women in science between PhD and faculty jobs^3^.

**Figure 1:**
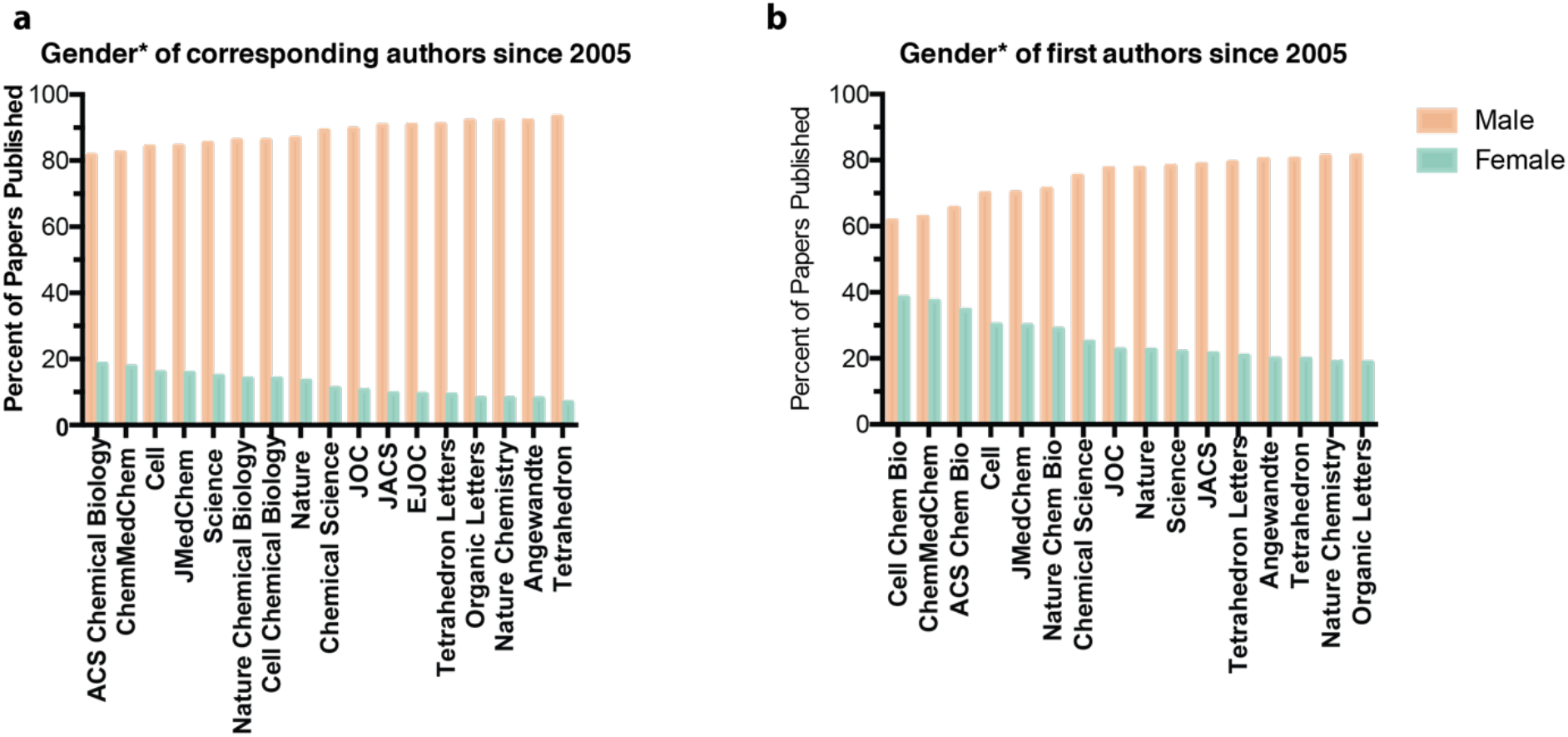
Rank-ordered gender breakdown of corresponding and first authors for papers published since 2005. A) Gender breakdown of corresponding authors for papers published in chemistry, chemical biology and NCS journals since 2005. B) Gender breakdown of first authors for papers published in chemistry, chemical biology and NCS journals since 2005. *Name-predicted gender–see methods section for details.

Next, we plotted the percentage of female and male corresponding authors for each journal since 2005. Given the results from the collated data we separated the journals into chemistry (figure 2a), medicinal chemistry (figure 2b), chemical biology (figure 2c) and NCS (figure 2d). Dashed lines on the graphs represent the US female and male academic faculty percentages in synthetic chemistry in 2020 (n = 518). There is a no apparent trend towards parity across all journals over the past 16 years. *Journal of Medicinal Chemistry* had the highest representation of women of all journals, with a 23.7% female corresponding author percentage so far for 2020.

**Figure 2:**
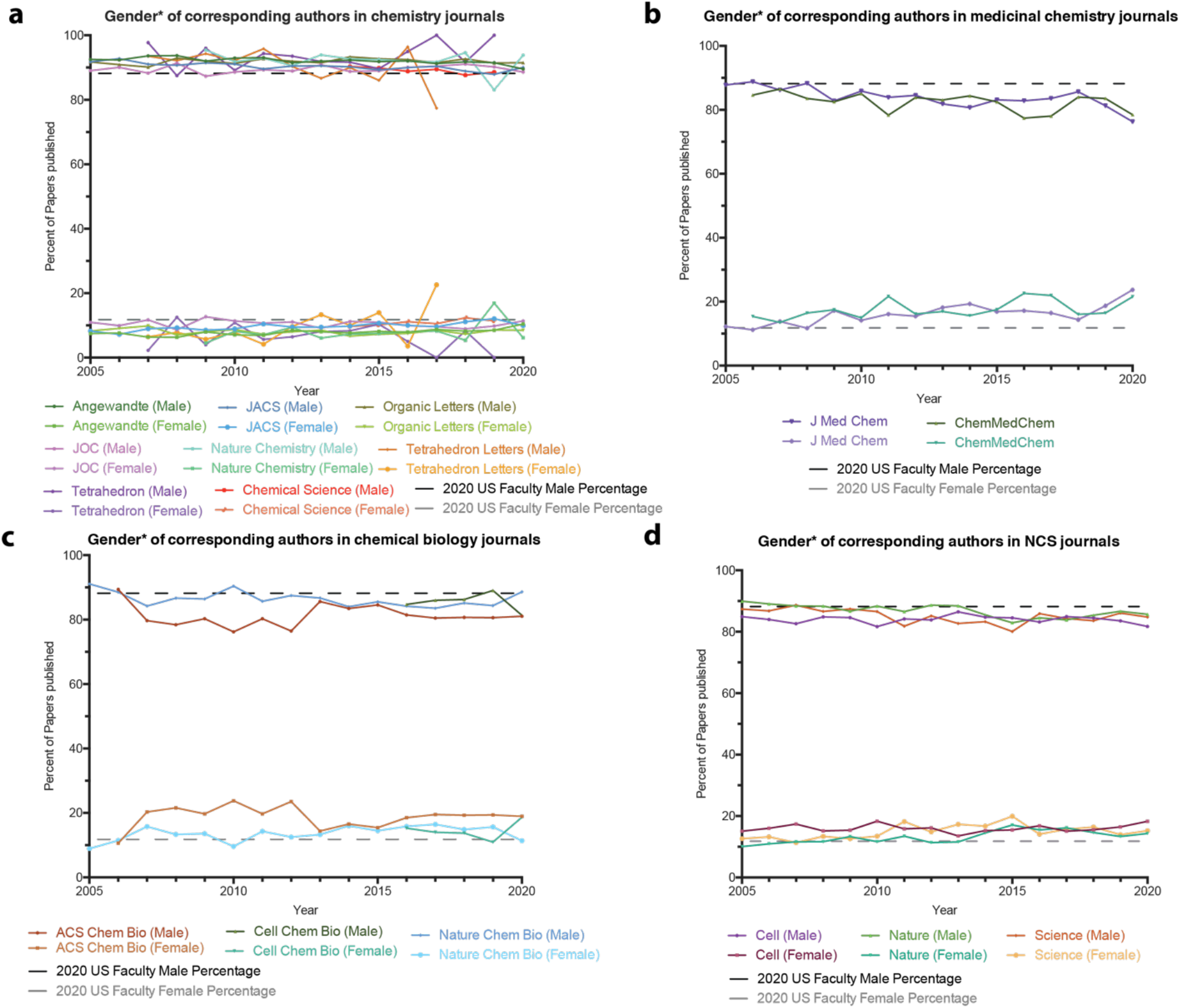
Gender percentages of corresponding authors for different journal types per year since 2005. Dashed lines represent current (2020) faculty percentages in US chemistry departments. A) Chemistry journals. B) Medicinal chemistry journals. C) Chemical biology journals. D) Nature, Cell, Science. *Name-predicted gender–see methods section for details.

We completed the same analysis as Figure 2, but for first authors. The dashed lines here represent NSF self-reported gender percentages for US graduate student and postdoctoral fellows combined. The gender percentages have stayed steady over the past 16 years (Figure 3). Similar to corresponding authors, underrepresentation of women is most pronounced in chemistry journals (Figure 3a). Both medicinal chemistry and chemical biology first author percentages have tracked closely with the percentages seen in graduate schools, suggesting women and men are equally likely to receive first authorship in these fields.

**Figure 3:**
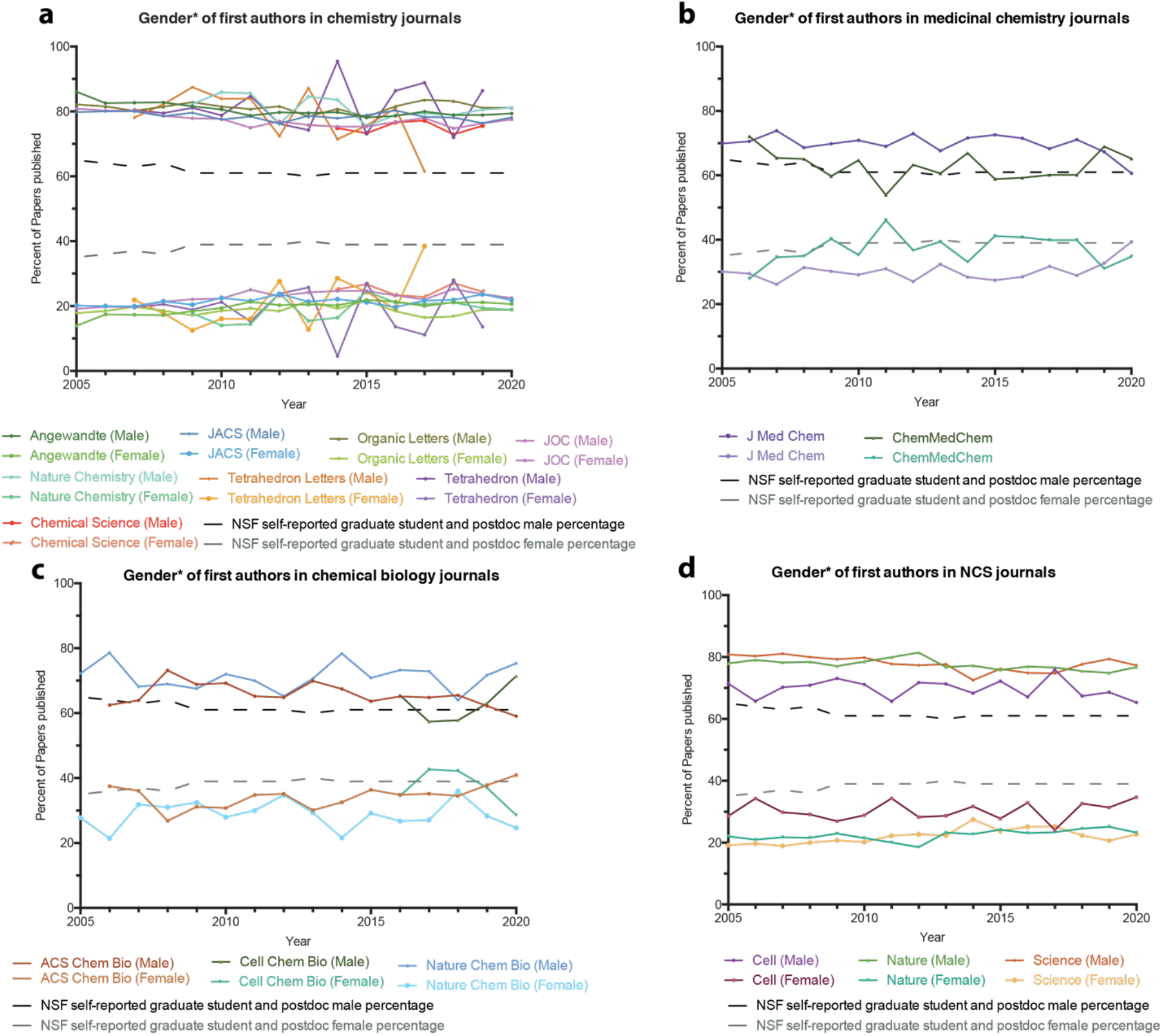
Gender percentages of first authors for different journal types per year since 2005. Dashed lines represent the NSF self-reported gender percentages of US graduate students and postdoctoral fellows. A) Chemistry journals. B) Medicinal chemistry journals. C) Chemical biology journals. D) Nature, Cell, Science. *Name-predicted gender–see methods section for details.

The final analysis we conducted on our data sets, was to assess if first authors were more likely to be women or men depending on the gender of the corresponding author. We show four representative journals in Figure 4, and the remaining data can be accessed in the Supporting Information. Across all journals first authors were more likely to be women if the corresponding author was a woman. In numerous cases, including all chemical biology journals, female corresponding authors had multiple years at 50:50 first author ratios. There is little to no trend in first author percentages over time, suggesting any progress towards parity in total first author ratios (Figure 3) may be due to the gradually increasing number of female faculty.

**Figure 4:**
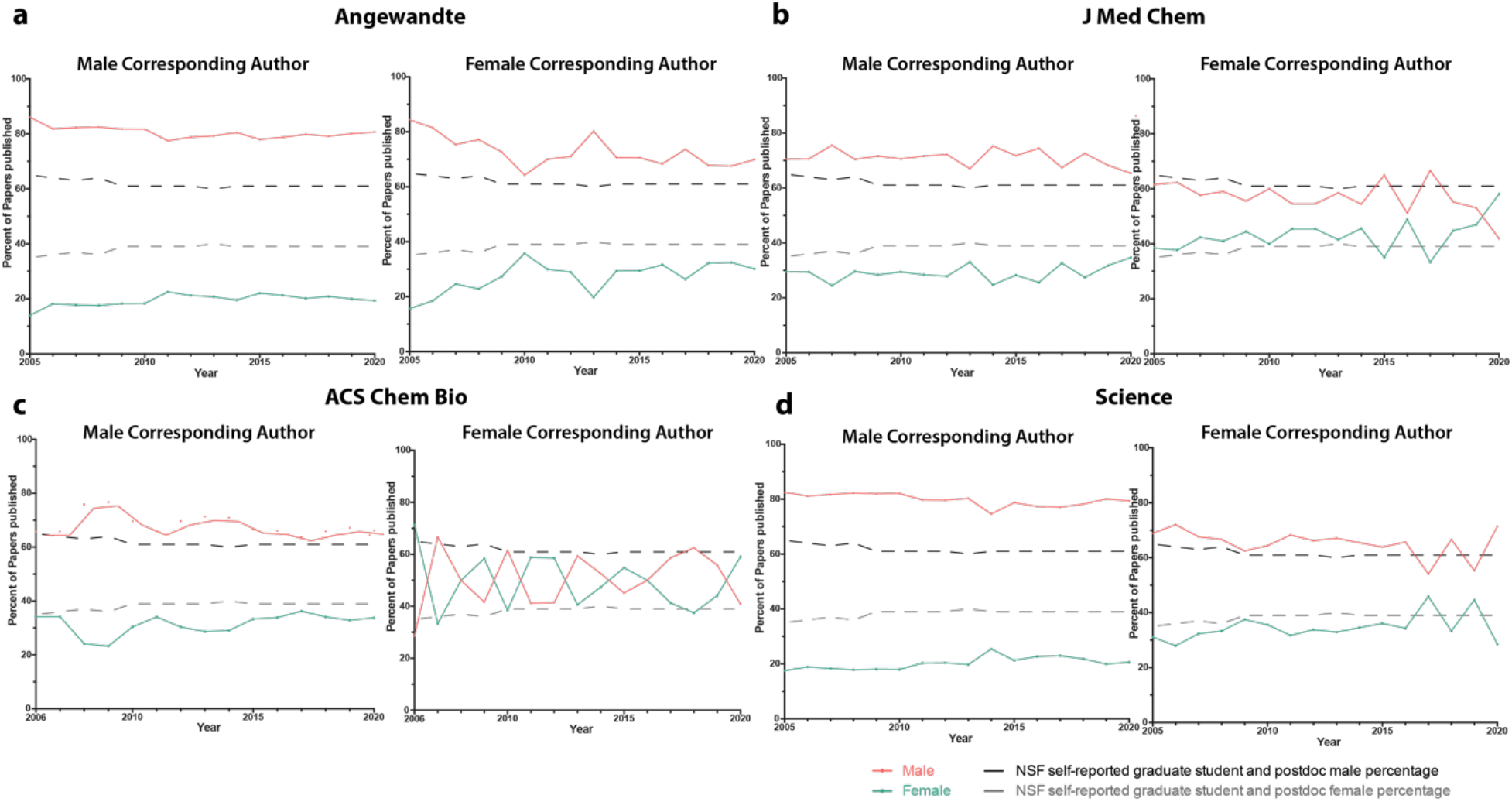
Gender^*^ percentages of first authors for both male and female corresponding authors. Dashed lines represent the NSF self-reported gender representation of US graduate students and postdoctoral fellows. a) First author gender percentage in Angewandte with either a male (left) or female (right) corresponding author. B) First author gender percentage in JMedChem with either a male (left) or female (right) corresponding author. C) First author gender percentage in ACS Chemical Biology with either a male (left) or female (right) corresponding author. D) First author gender percentage in Science with either a male (left) or female (right) corresponding author. *Name-predicted gender–see methods section for details.

In addition to authorship data, we were interested in gender representation on journal editorial boards. We manually curated names and countries of editorial board members from journal websites and used the same protocol with Gender API to estimate gender percentages. Due to the wide breadth of topics covered in Nature, Cell, and Science, the editorial boards for these journals were not analyzed. Editorial board gender ratios ranged from ~1:9 to 1:1 female: male. (Figure 5). Tetrahedron and Tetrahedron Letters have journal advisory boards of over sixty members in which less than 10% of the members are women. The boards of Nature Chemistry and Nature Chemical Biology are 50% and 60% female, but it should be noted that these journals differ from the others in that they have a small team of full-time editors.

**Figure 5:**
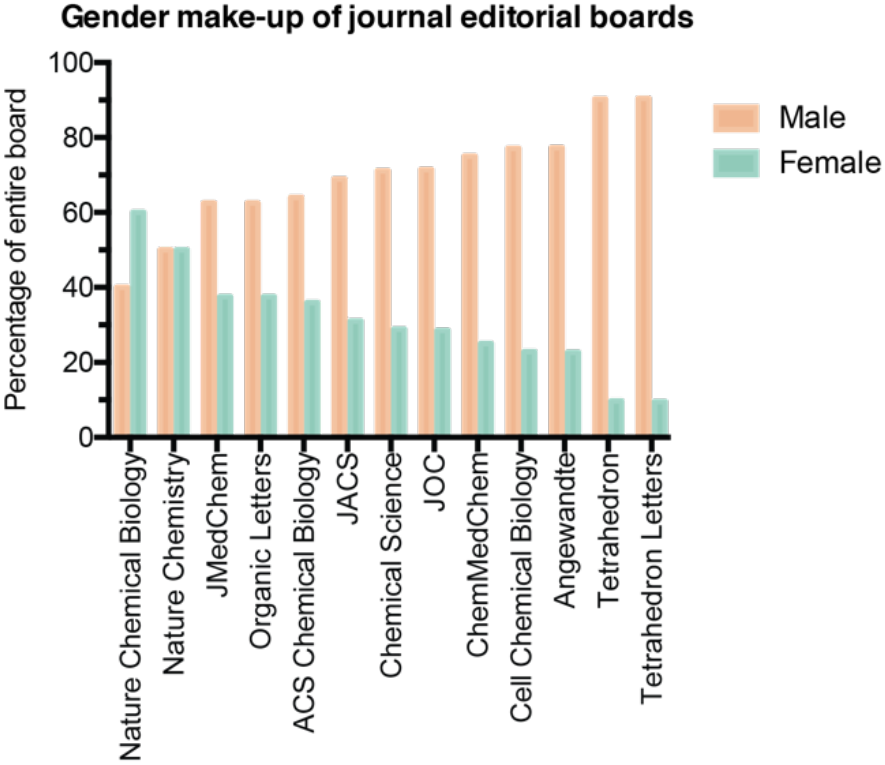
Rank-ordered gender ratios of journal editorial boards in 2020. Data collected from respective journal websites.

## Conclusion

Some salient trends emerged in our dataset:

1. Chemistry journals have a large disparity in gender ratios, especially for corresponding authors.
2. ACS Chemical Biology was the journal that with female to male ratios closest to parity, at 35:65 and 18:82 for first and corresponding authors respectively.
3. In the past 16 years there has been no increase in the representation of women as either corresponding or first authors.
4. There is a large drop off in female representation between graduate school and faculty positions.
5. First author representation is more likely to be equitable when the corresponding author is female.
6. With few exceptions, the gender ratios of editorial boards heavily favor men.

It is clear that, despite recent publications on the topic, there is still a large problem in our field that needs to be addressed. The goal of our analyses is to provide data on authorship which we hope will foster further discussion about how to improve gender representation in chemistry. Open discussion on the topic of diversity yields creative ideas and we hope our data allows informed conversations about how we as a community can improve representation of women in chemistry and the wider sciences. Additionally, it is our hope that this work can evolve to be more inclusive, and will promote discussion about other marginalized groups.

## Methods

On June 10^th^ 2020 a web-scraper was used to search PubMed for all papers published in a journal each year since 2005. For journals that were created post-2005, their date of creation was used as the first year. The scraper pulled the first name of both the first and corresponding authors. The country of the corresponding author was extracted, and this was assumed to be the same for the first author. The country information improves the accuracy of the gender predicting algorithm. Any papers that have a solo author were discarded from the data. This resulted in 196,157 articles. The data sets containing a first name and country were uploaded to Gender API to predict the gender of the author based on their first name. Gender API outputs a predicted gender, an accuracy score out of 100 and data size sampled. The accuracy score is a probability of the gender being correctly assigned, thus any data points below 95% confidence or small sample size (below 100, with confidence below 100) were removed from the data set. This resulted with a set of high-quality data points for 296,245 authors. A histogram demonstrating the quality of the data set is found in supporting information. Any years for a journal with less than 25 quality data points was removed from any time-series analysis. Each author was then assigned a binary gender and population totals calculated.

NSF self-reported data were used to calculate the US graduate student and postdoc gender percentages. These data sets are available for 2005-2014; post-2014 values were extrapolated from the publicly available data.

ACS Division of Organic Chemistry has a curated list of 518 current US-based faculty who work on synthetic organic chemistry. Similar to above, a web-scraper was used to extract first-names, and Gender API was used to assign a name-predicted gender.

Journal websites were used to obtain their editorial boards. These lists were manually extracted, and board members’ genders manually assigned based on lab websites and publicly available data.

Our approach has some inherent limitations. While the use of a gender API is an efficient high throughput method to attempt to address gender prediction, gender is not conclusively discernible by name, so this approach is inherently reductionist^4^. Furthermore, due to the limitations of first name-based gender APIs, we could only account for two genders: male and female. We desired to include other genders in the analysis; however, we could not find a method to include them that was high throughput. We are aware that only considering male and female as genders can be considered an act of erasure of other genders, and as more data becomes available, we will include nonbinary genders to remedy this. Our approach does not account for gender transition, differing gender identity and gender expression, or nonbinary identities, which can give rise to several additional dynamics that are not captured with our data or discussion.

Additionally, we used the affiliated country of the last (corresponding) author to inform gender assignment by the API, which uses country to improve gender prediction for common names. There are likely cases in which that affiliated country will not reflect the origins of the names of the first or last authors. Additionally, there are likely cases where authors intentionally use a gender-neutral or male first name or publish only under their initials to avoid publishing bias. Finally, there are many cases where there are multiple first or multiple corresponding authors, and, more rarely, cases where the last author is not a corresponding author. Our approach does not account for these cases. Gender neutral names were also eliminated from the data sets in order to minimize uncertainty.

## Supporting information

Journal percentages tabulated

Corresponding author percentages tabulated

## Acknowledgements

We thank Etan Schwartz for help with analyzing the data sets. We thank James Fraser and Seemay Chou for helpful discussion. Iris Young and Roberto Efrain Diaz provided extremely helpful feedback on the first published draft of the manuscript

