## Supporting information for "Examining gender imbalance in chemistry authorship"

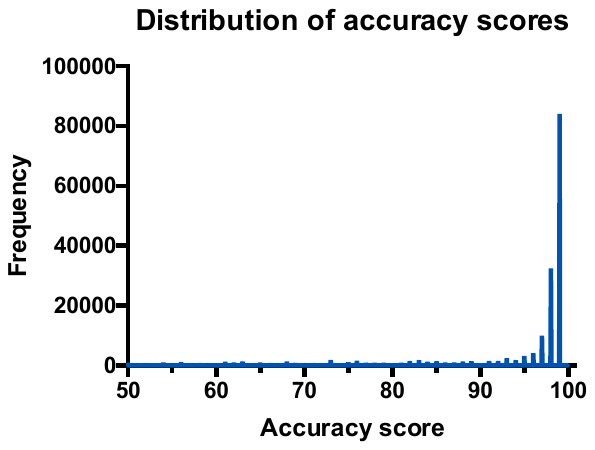
Histogram of accuracy scores

Corresponding author percentages by journal

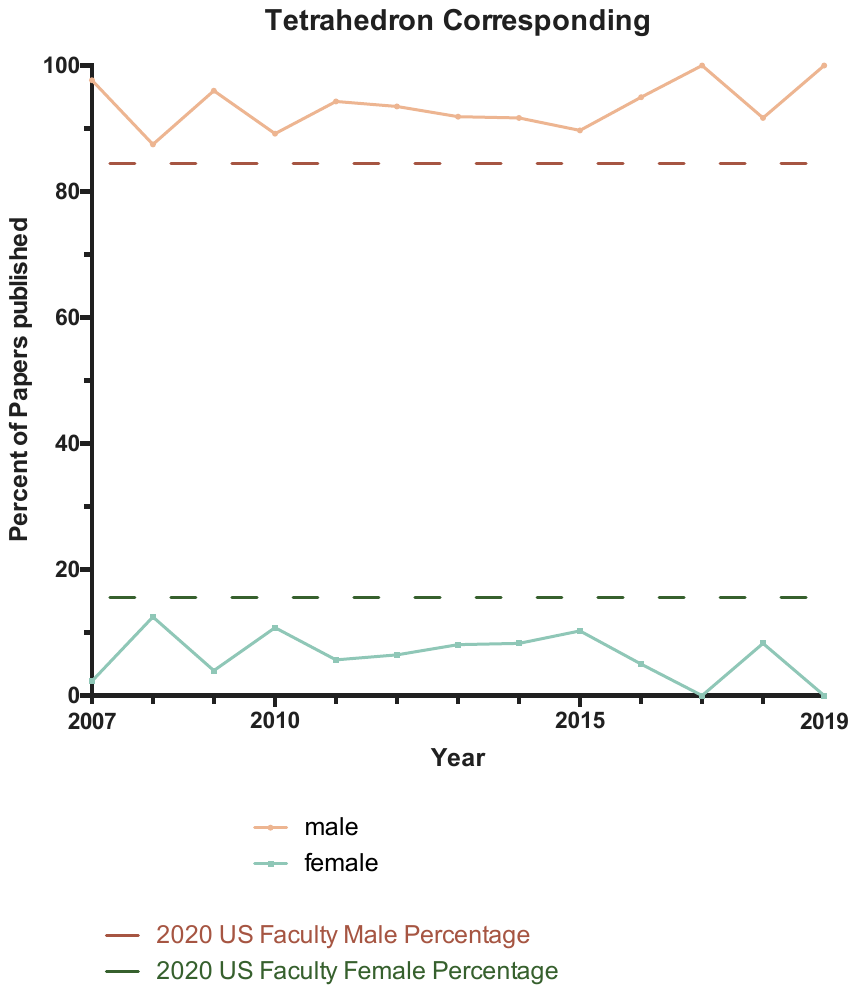

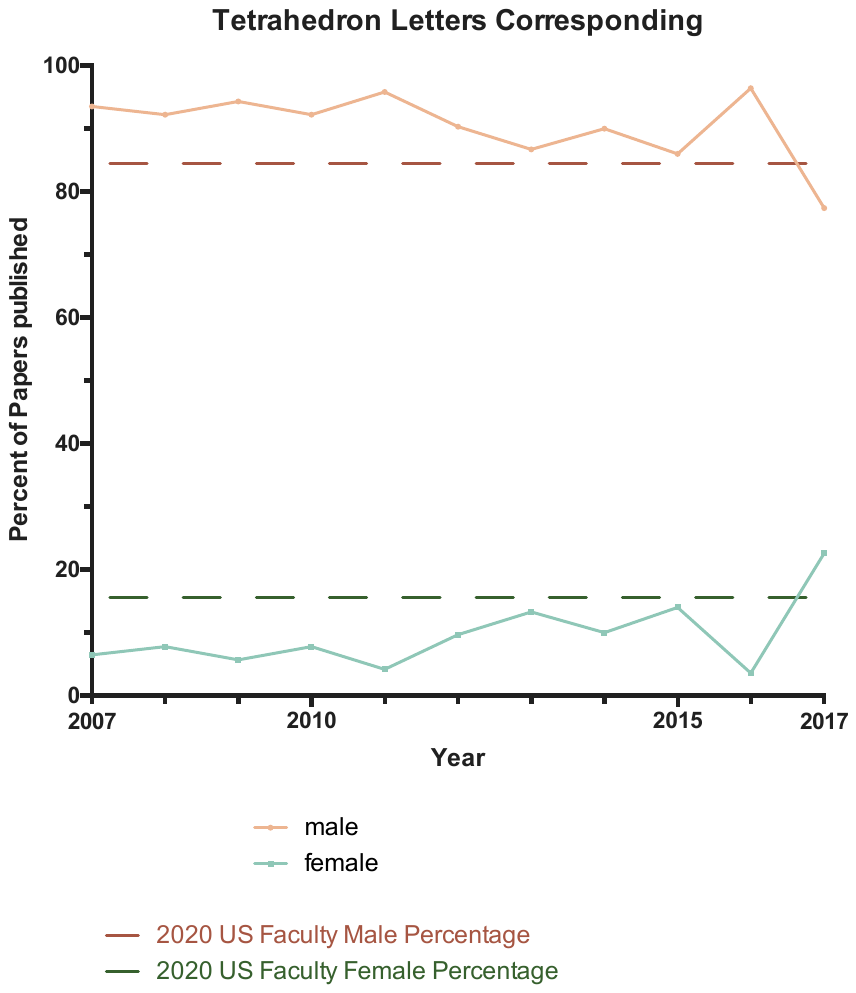

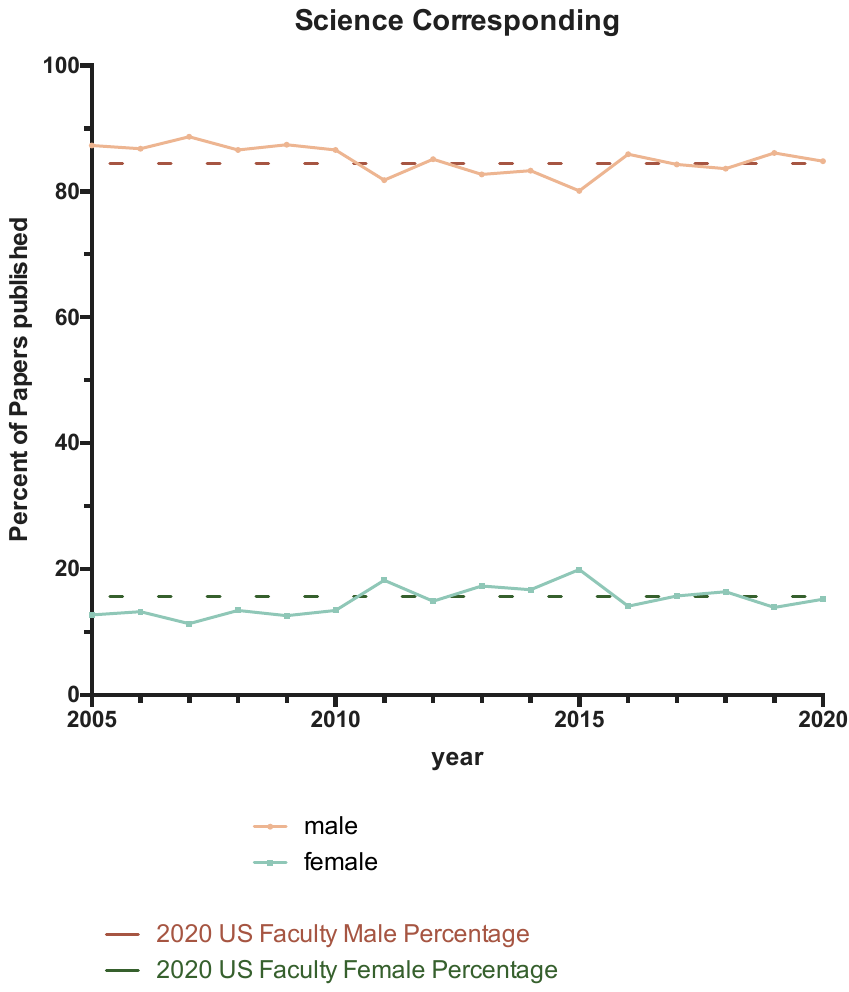

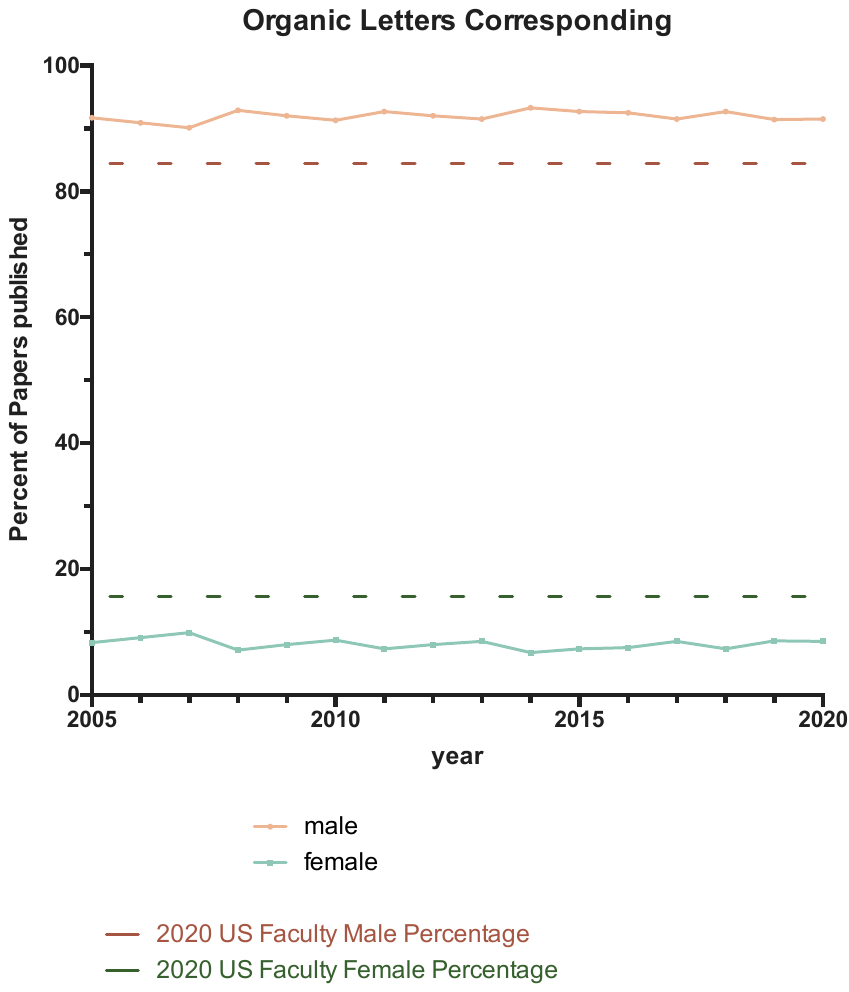

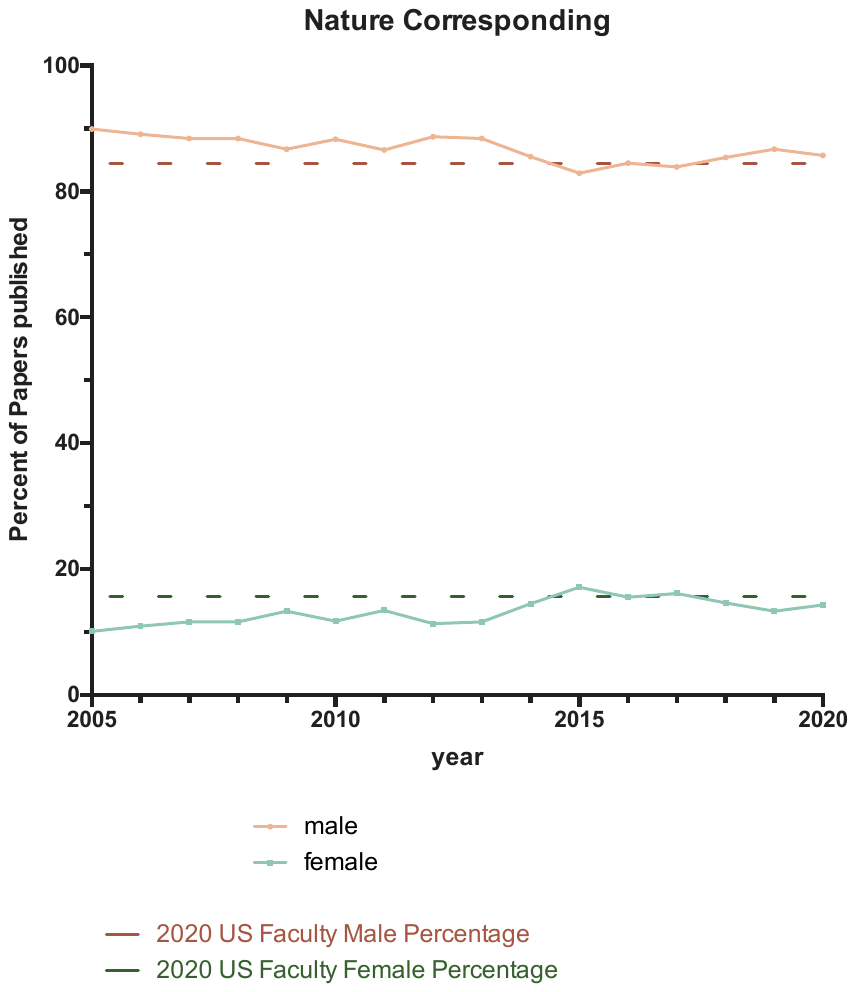

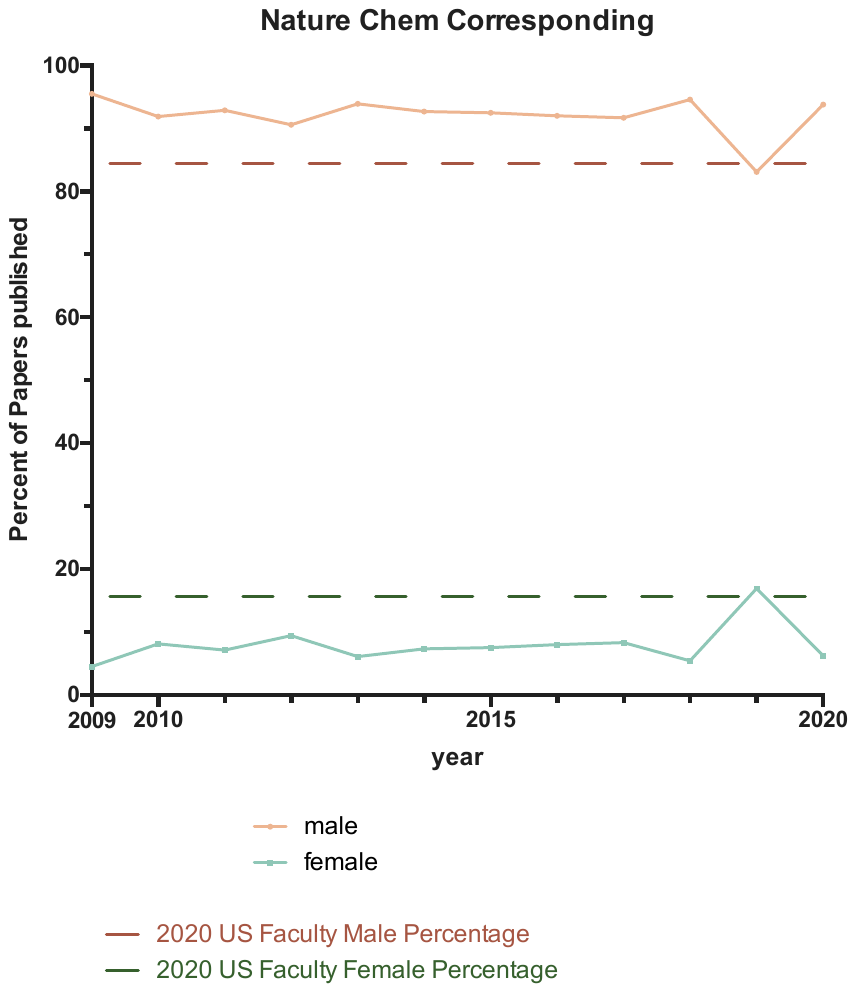

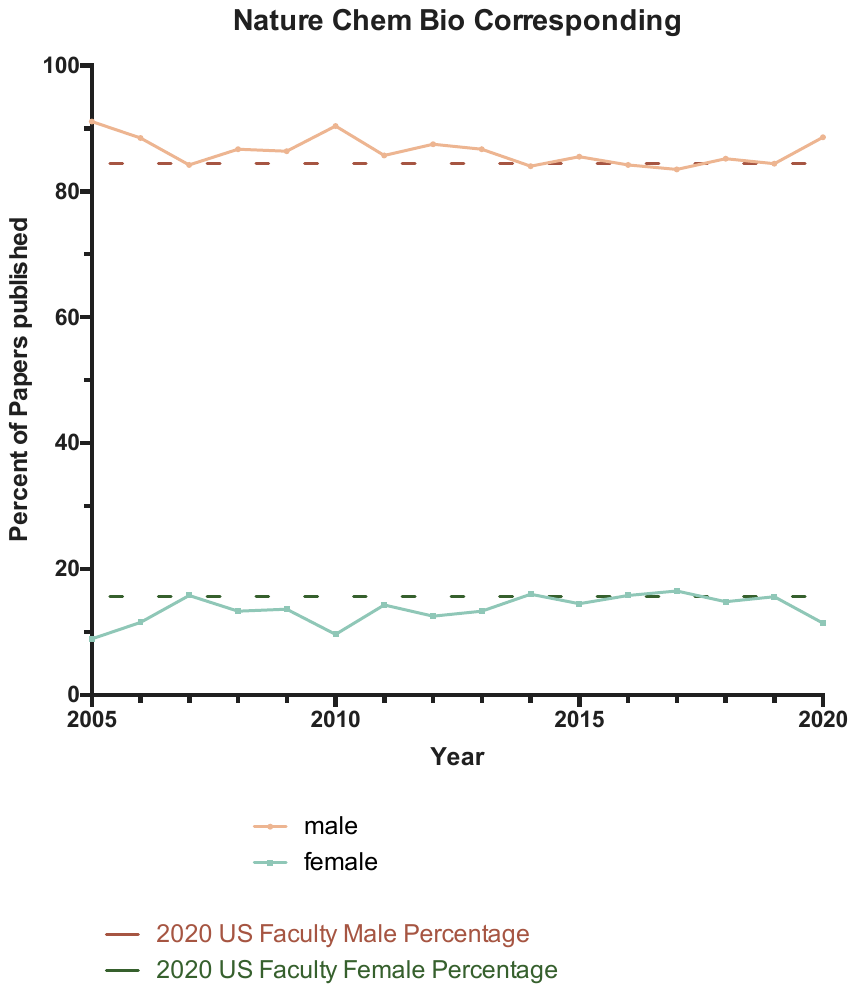

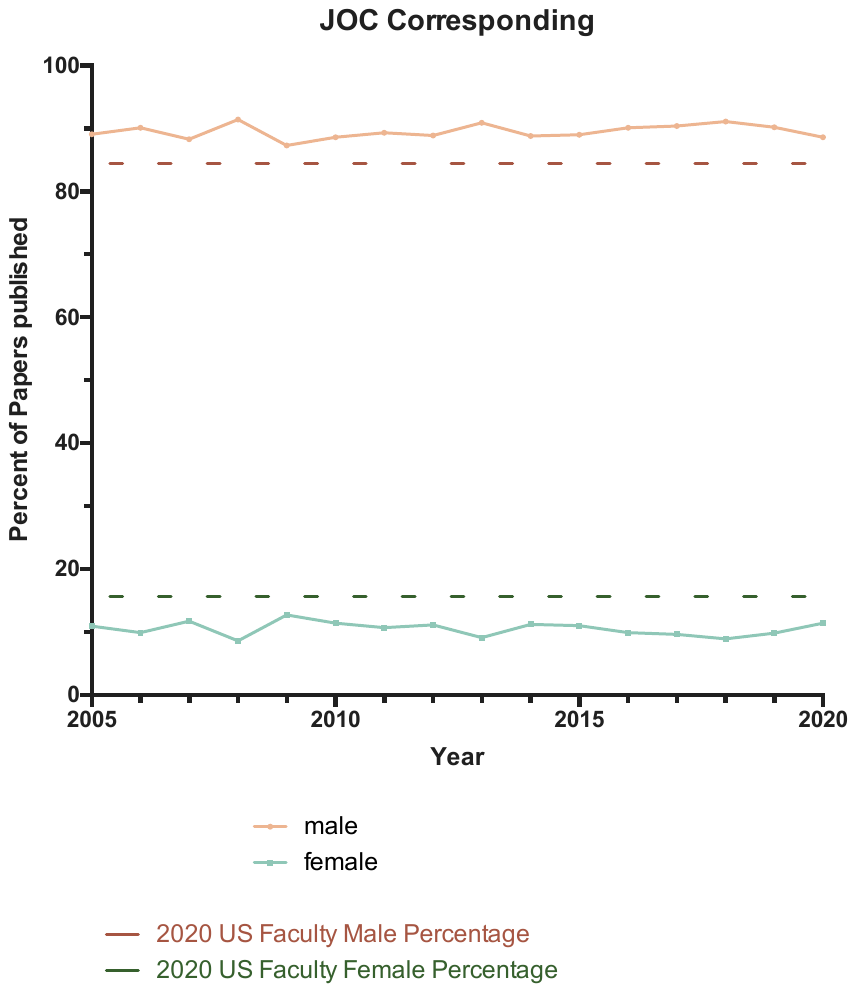

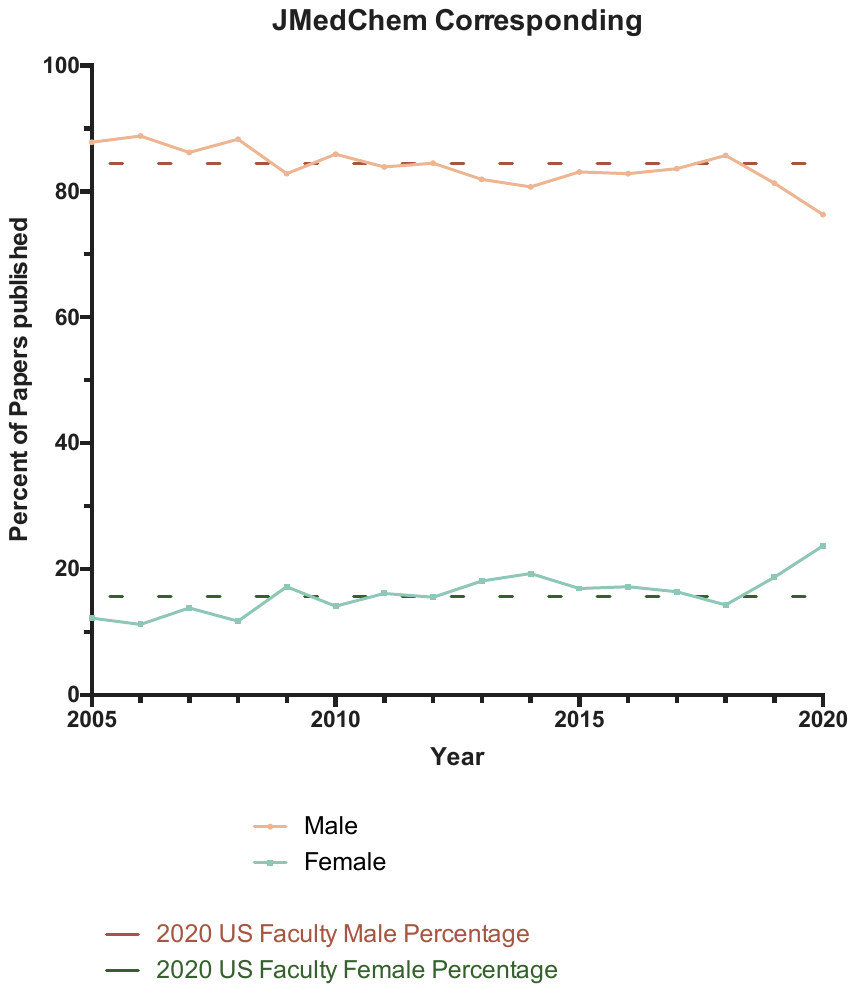

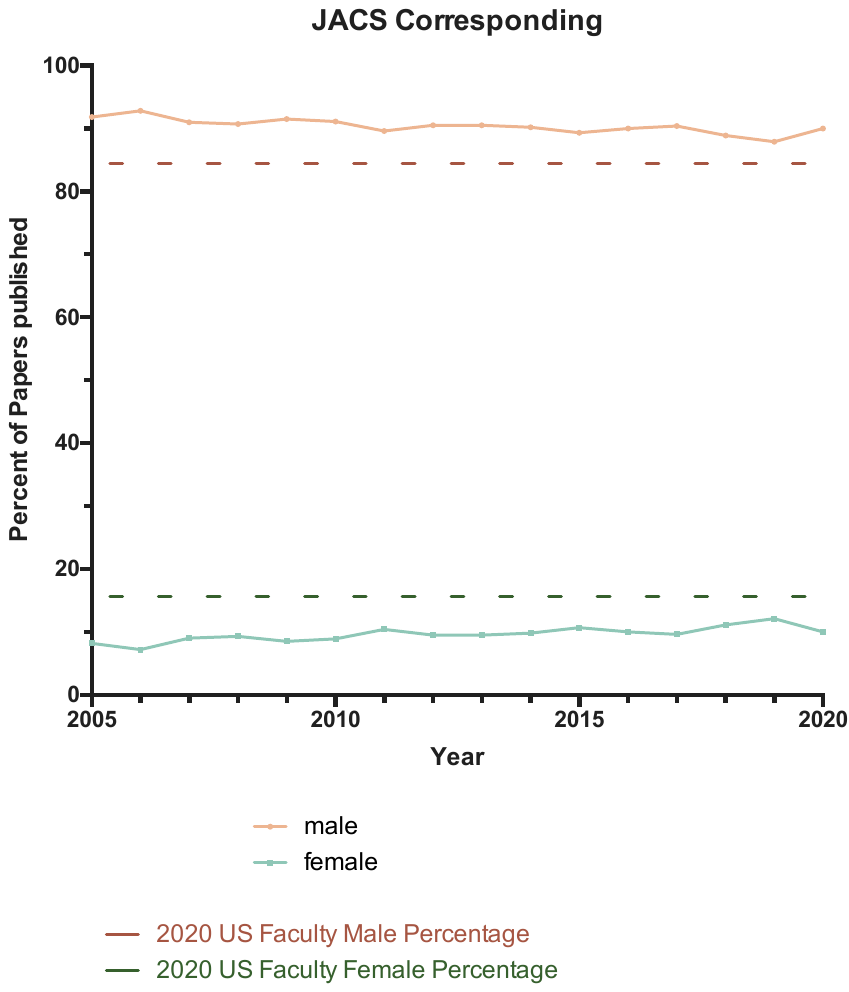

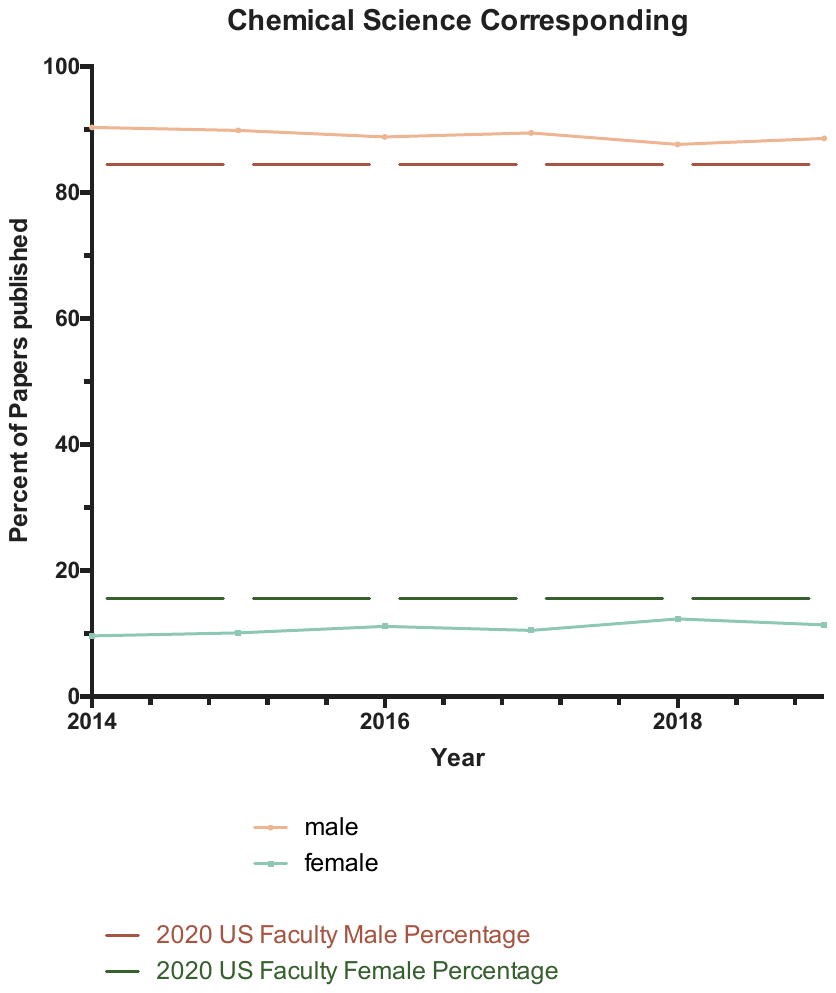

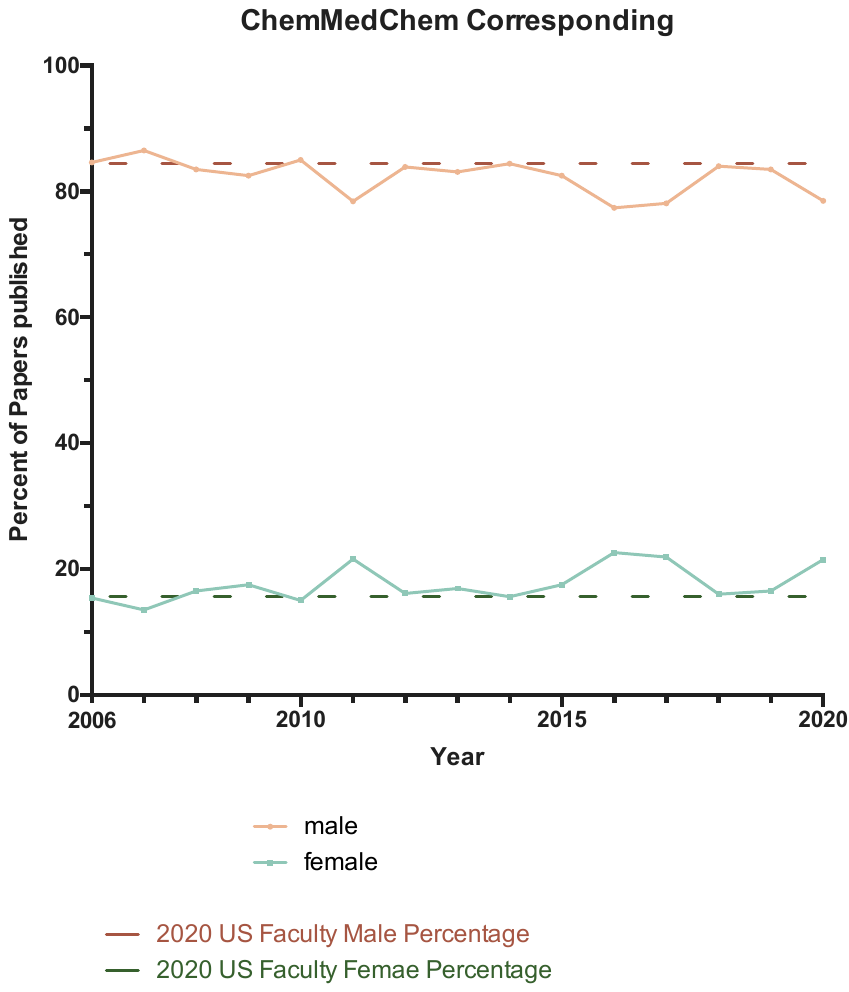

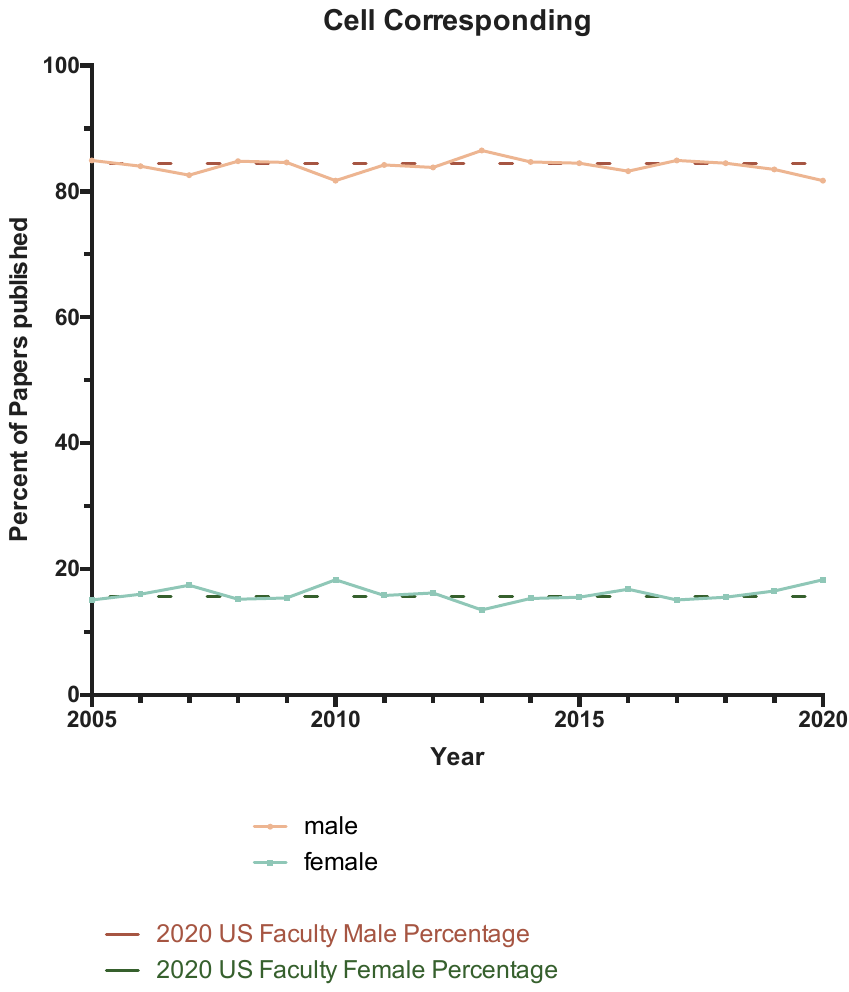

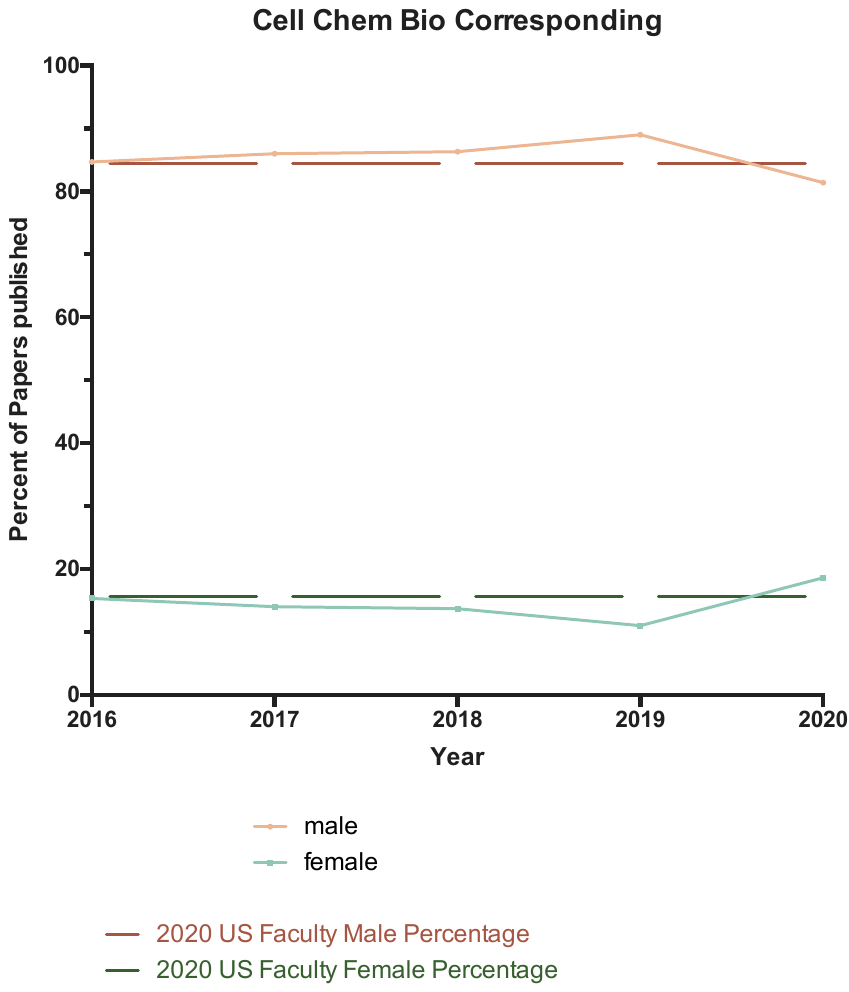

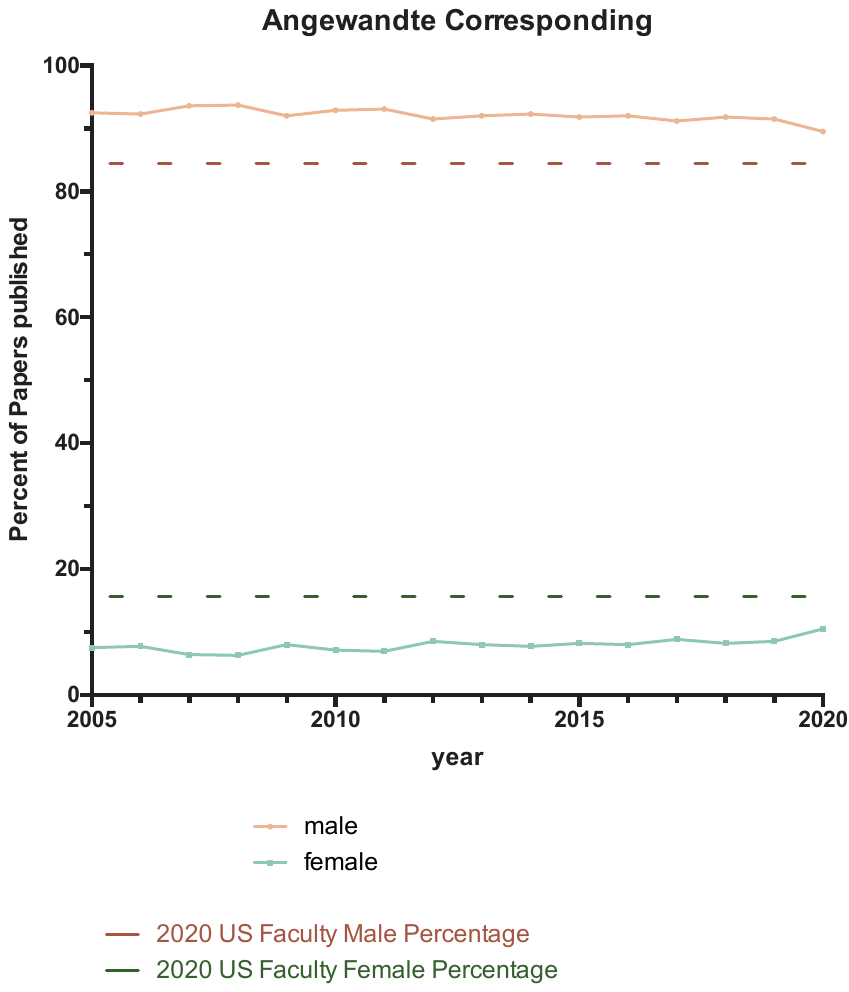

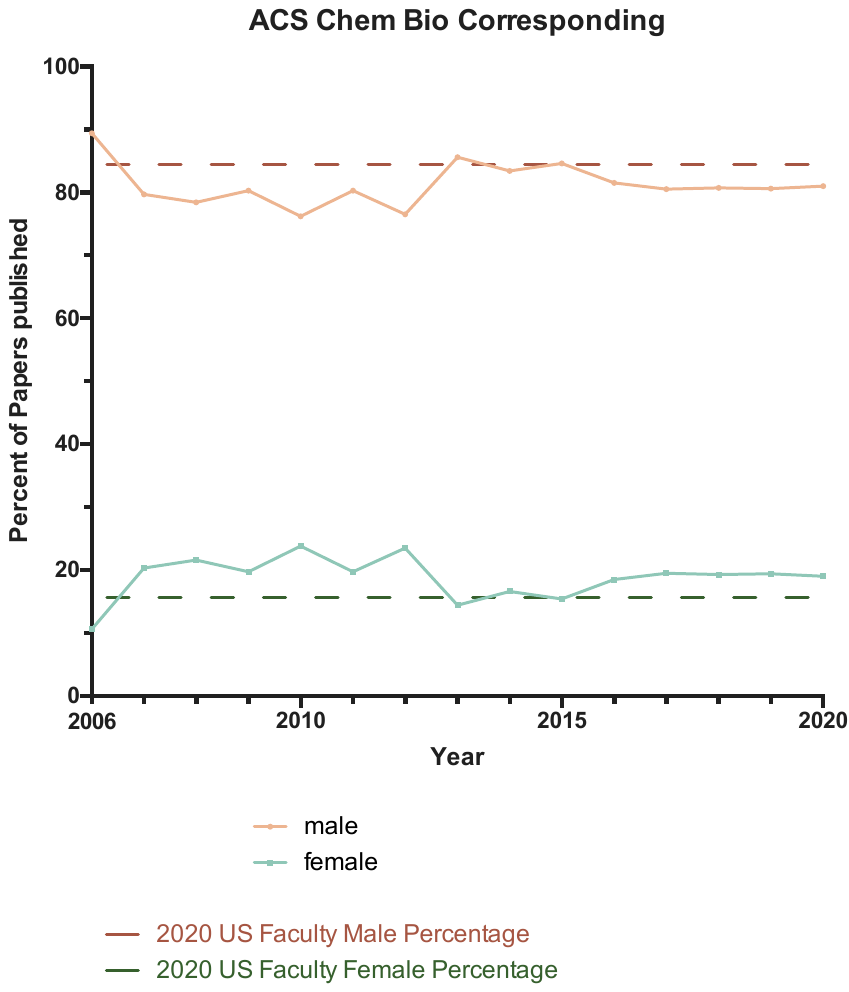

First author percentages by journal

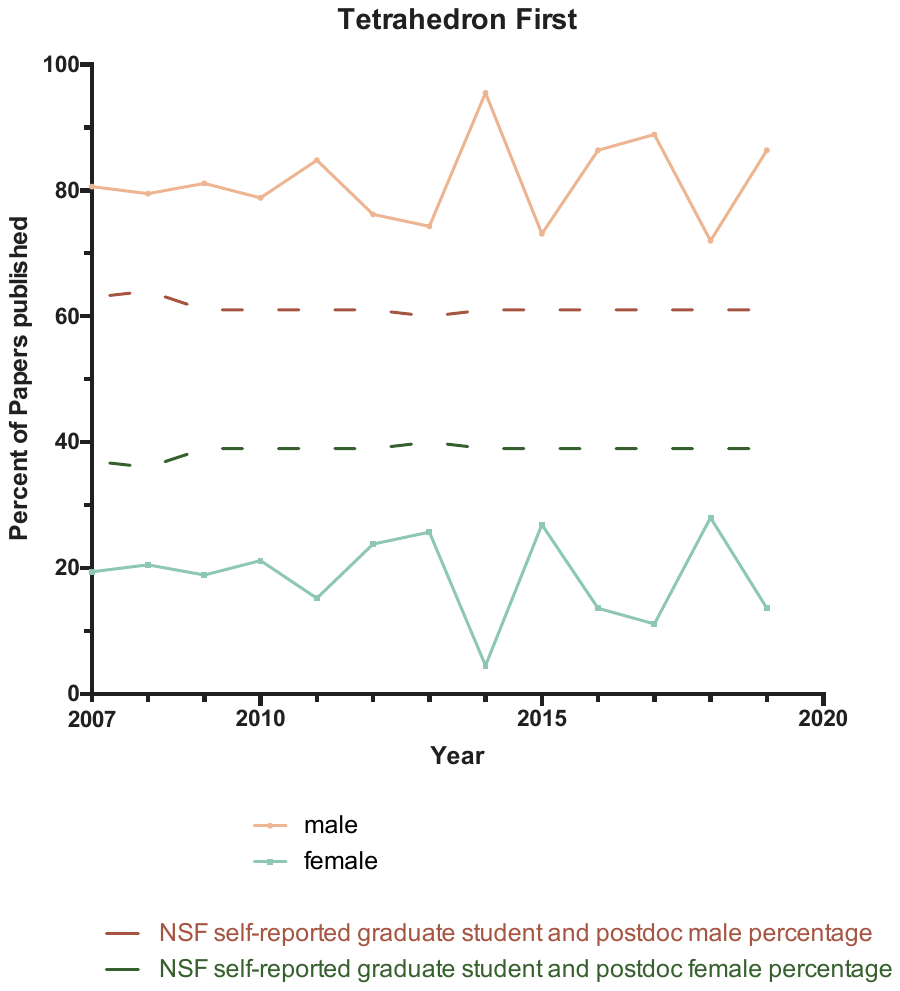

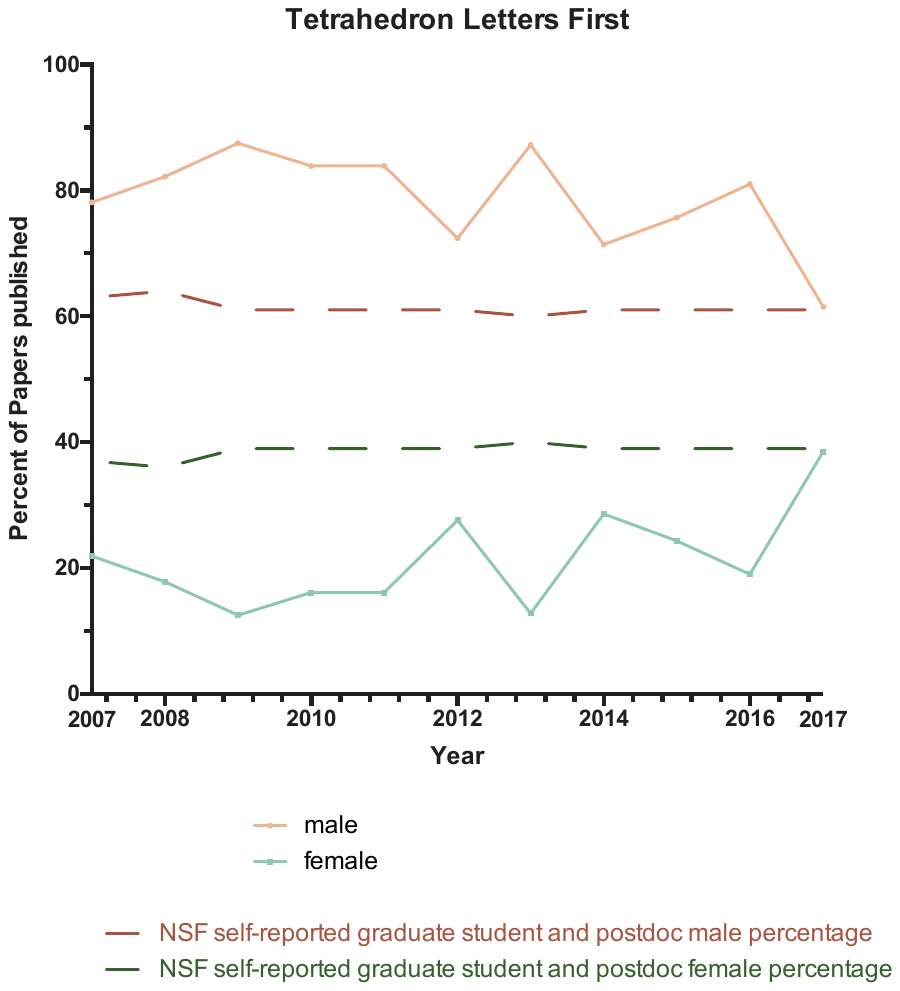

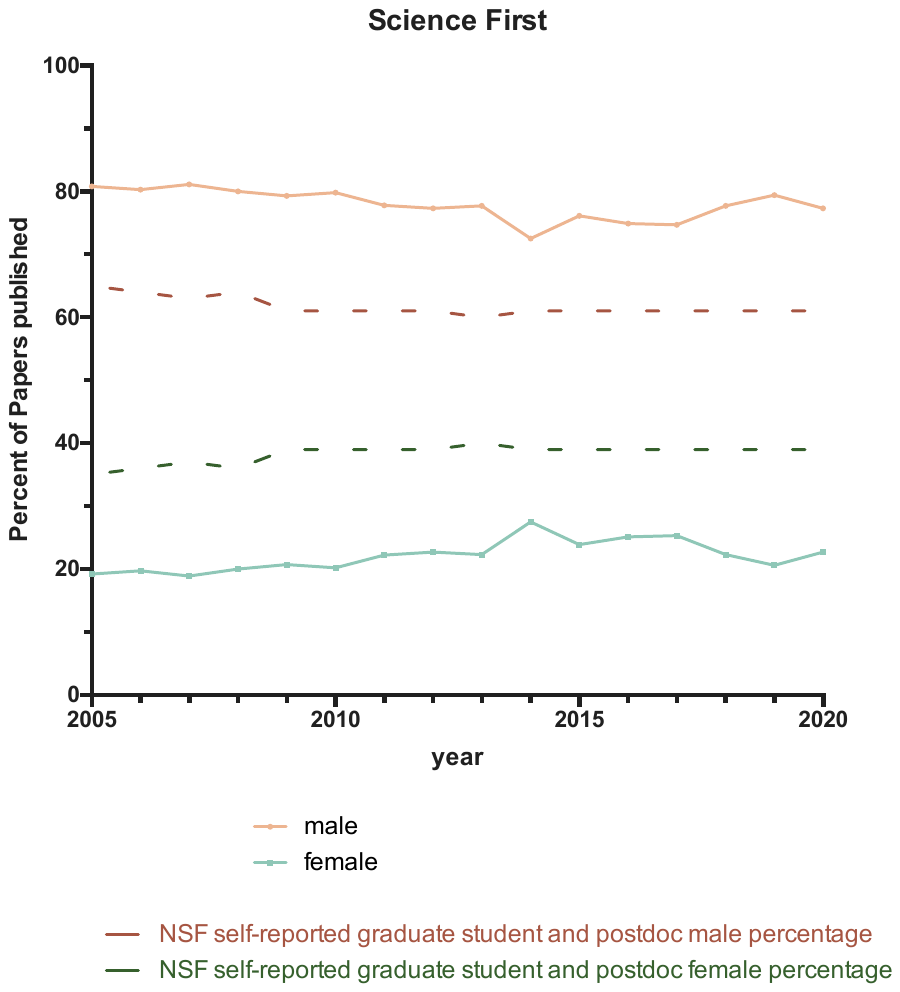

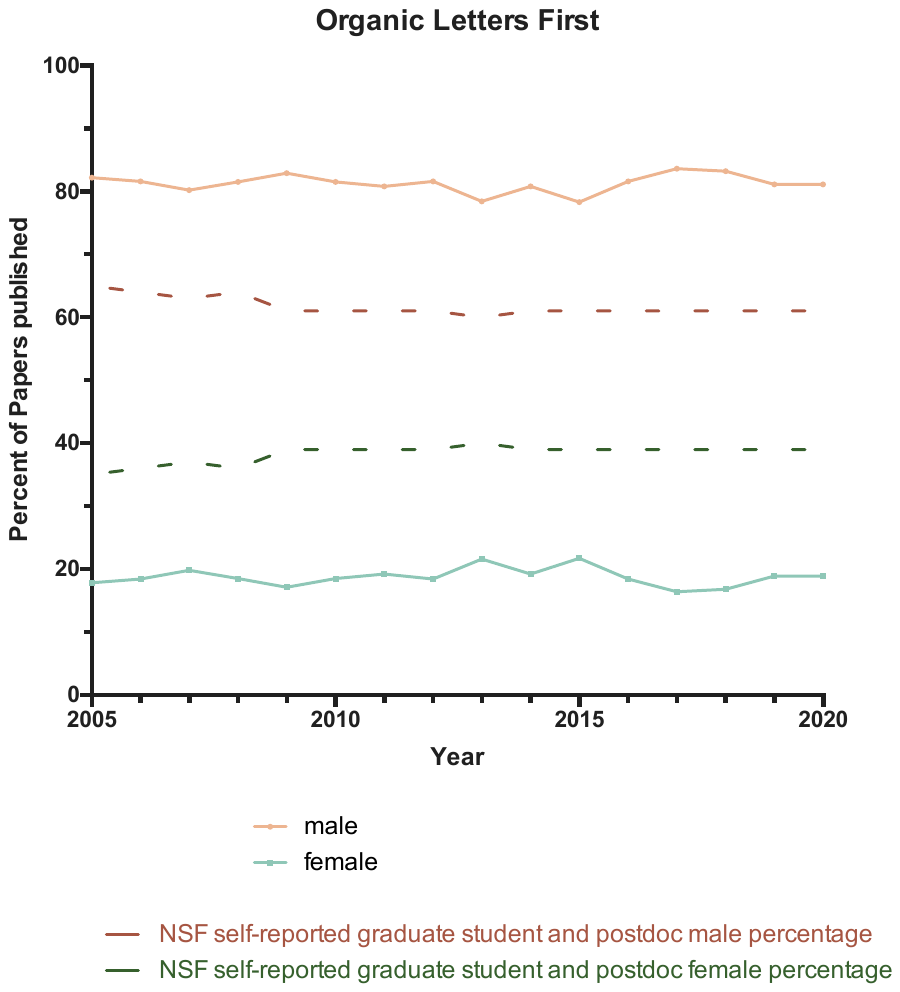

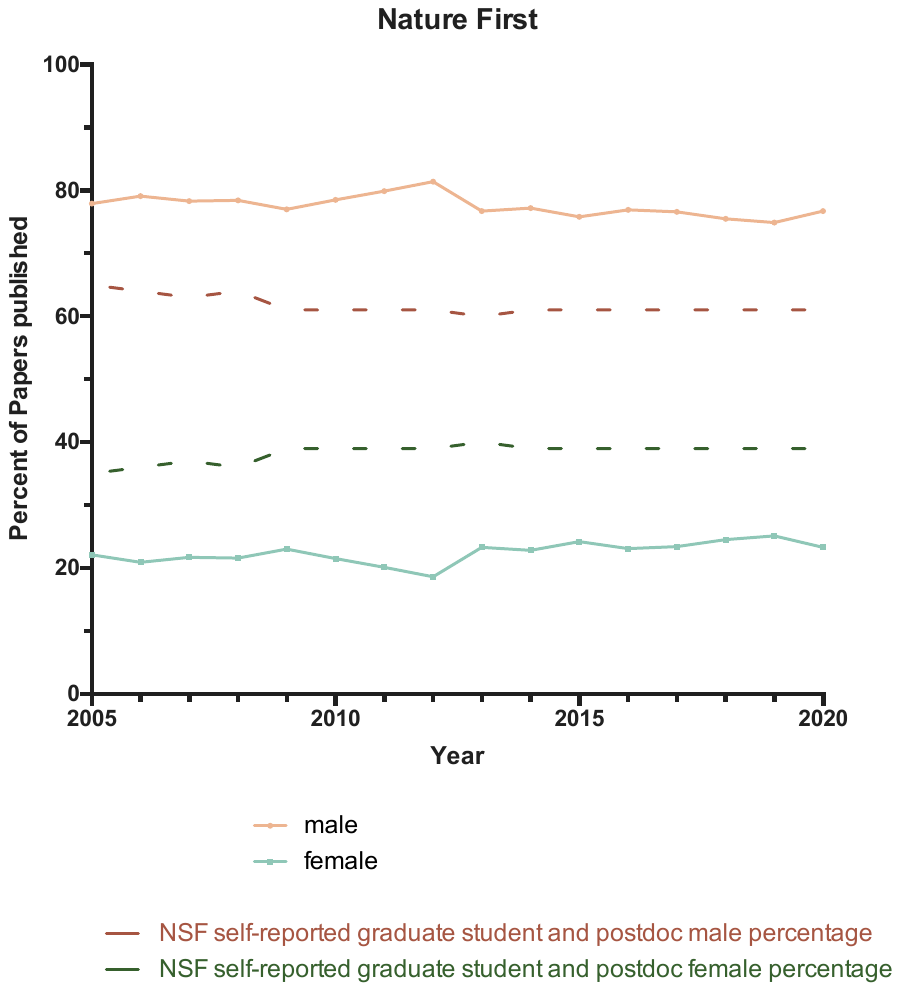

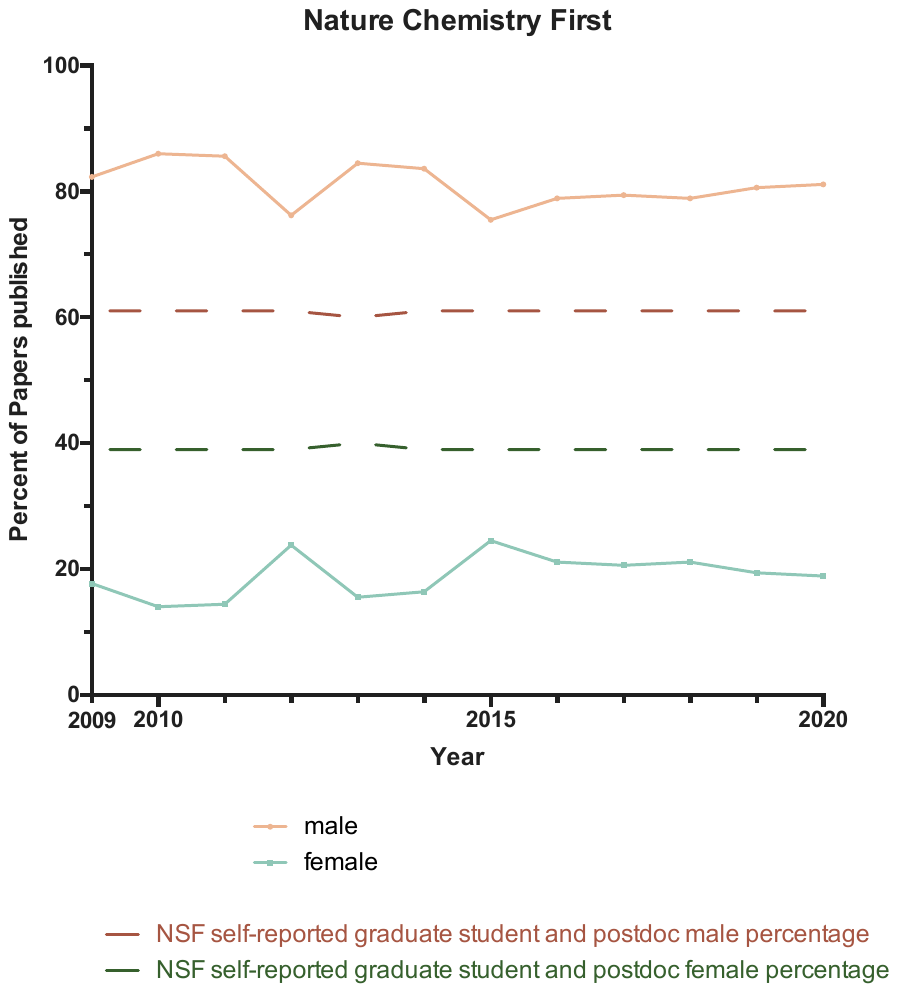

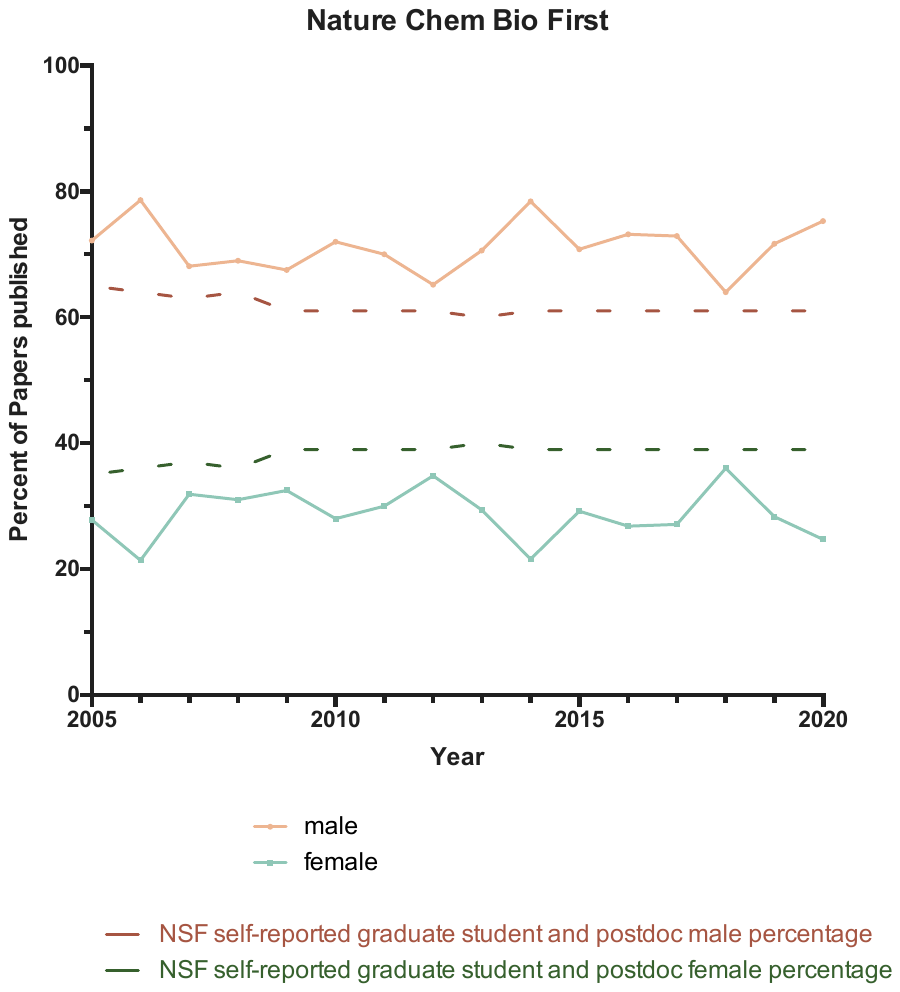

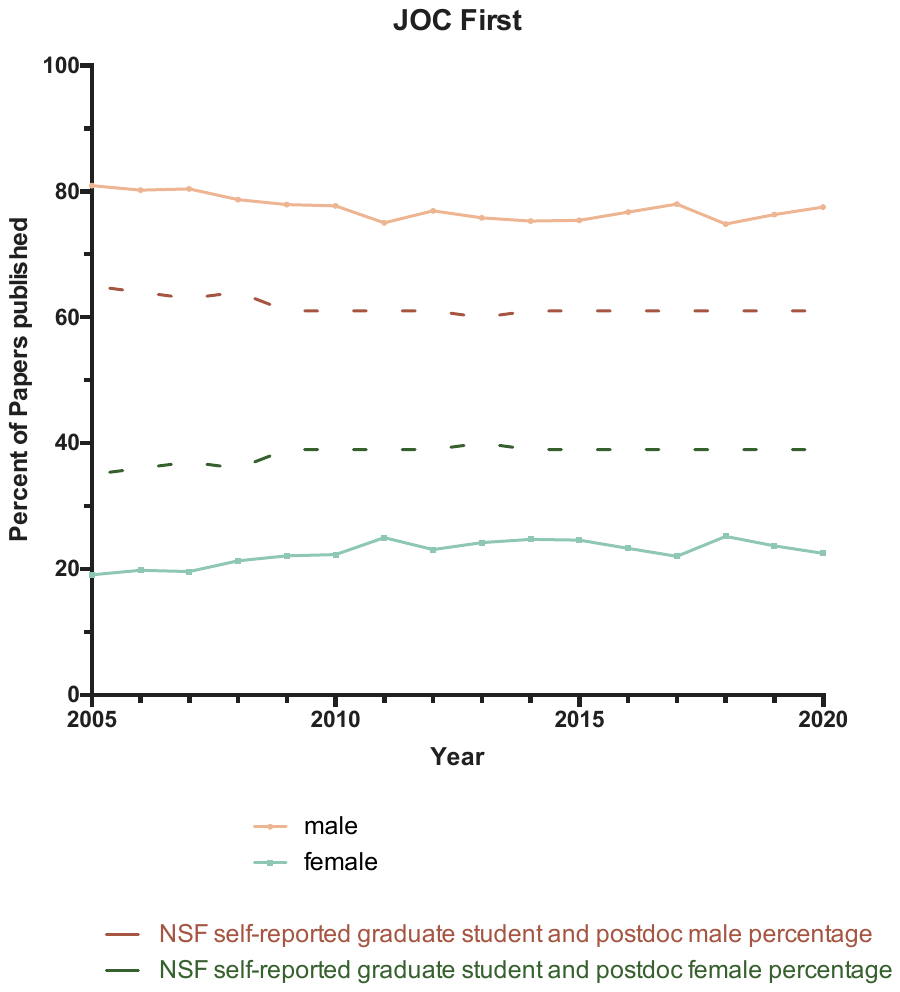

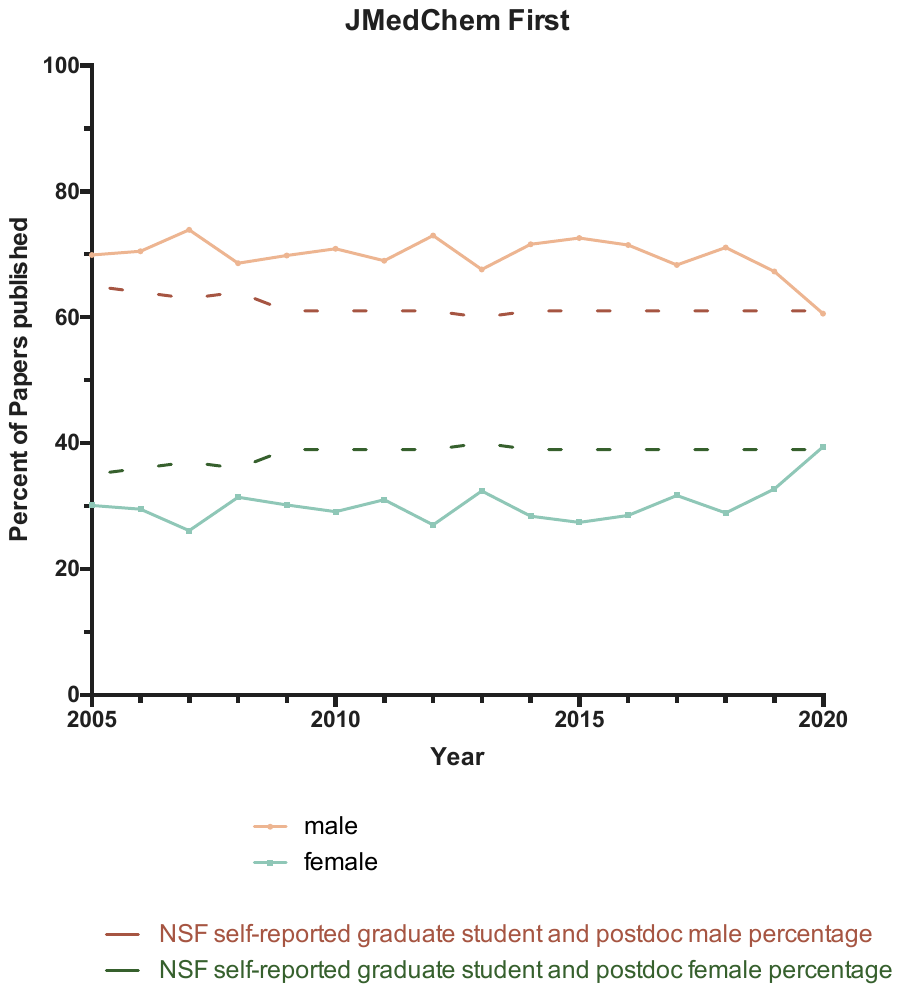

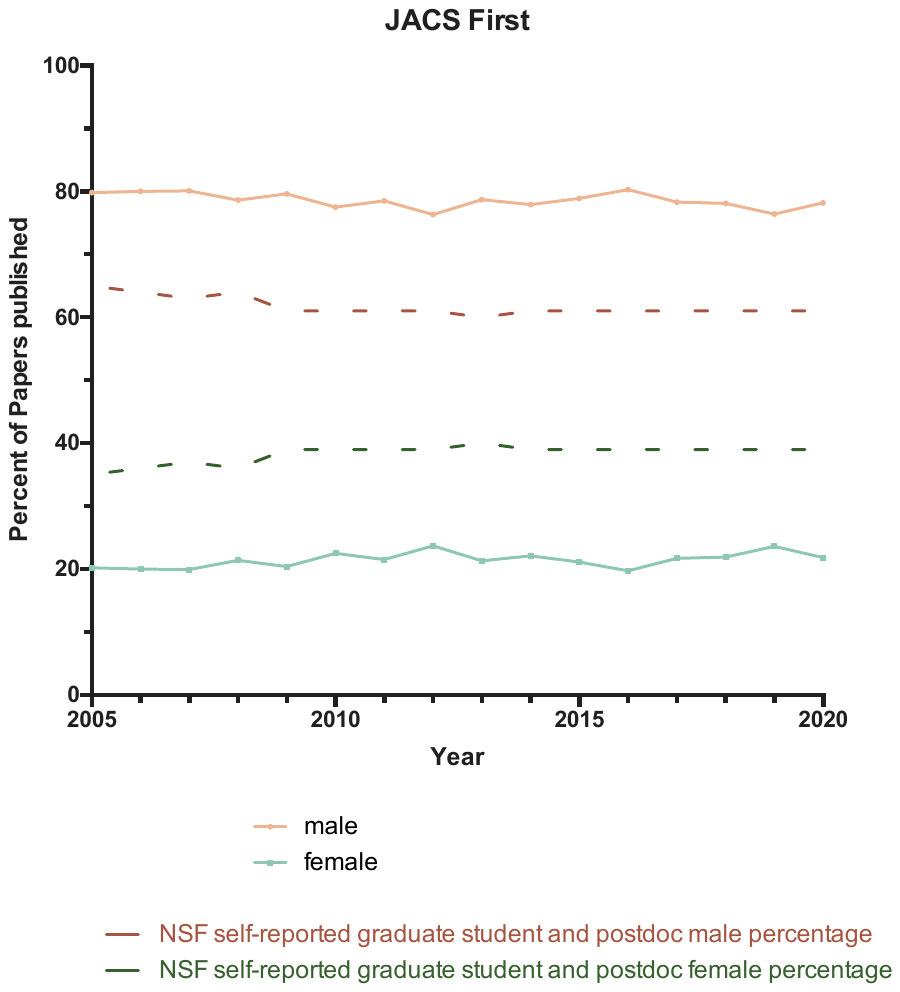

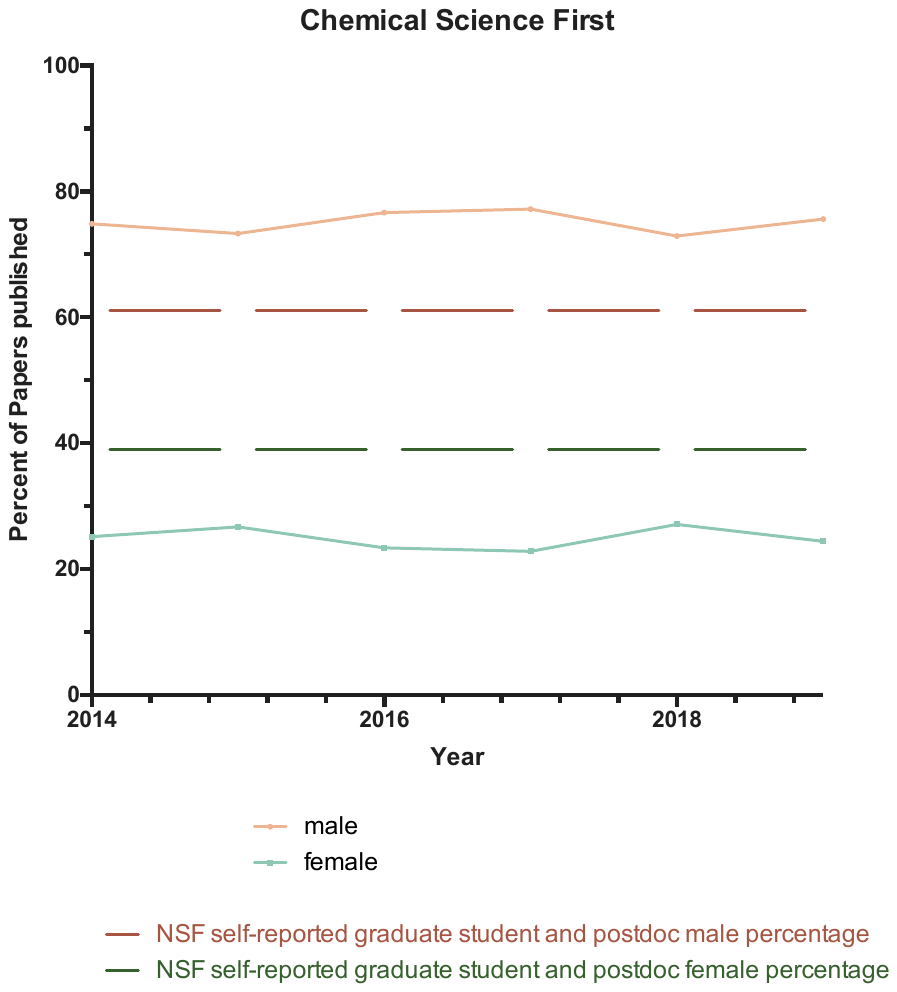

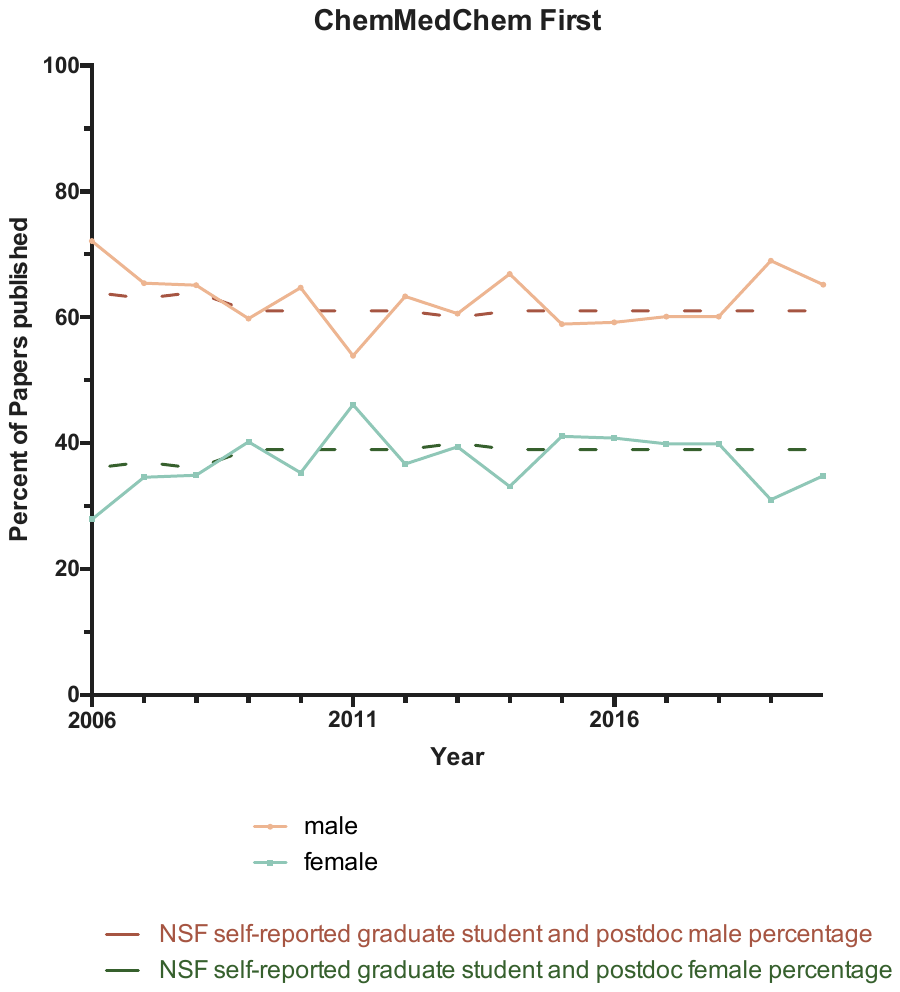

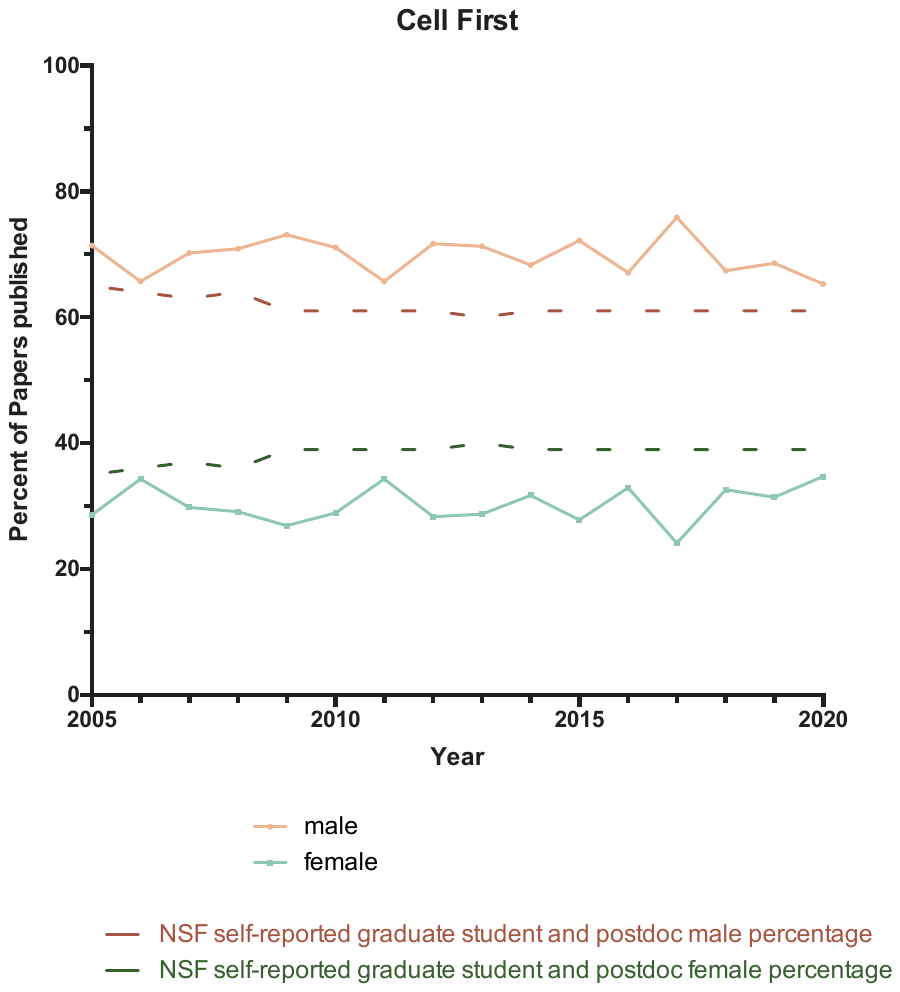

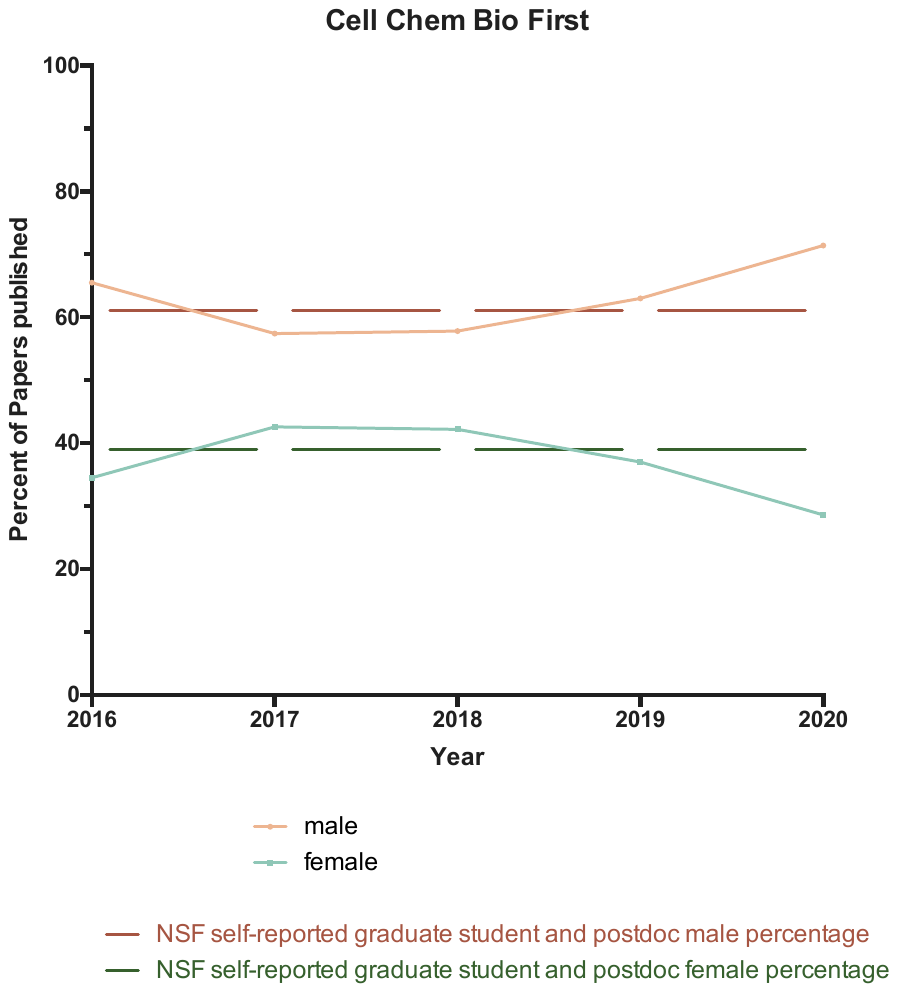

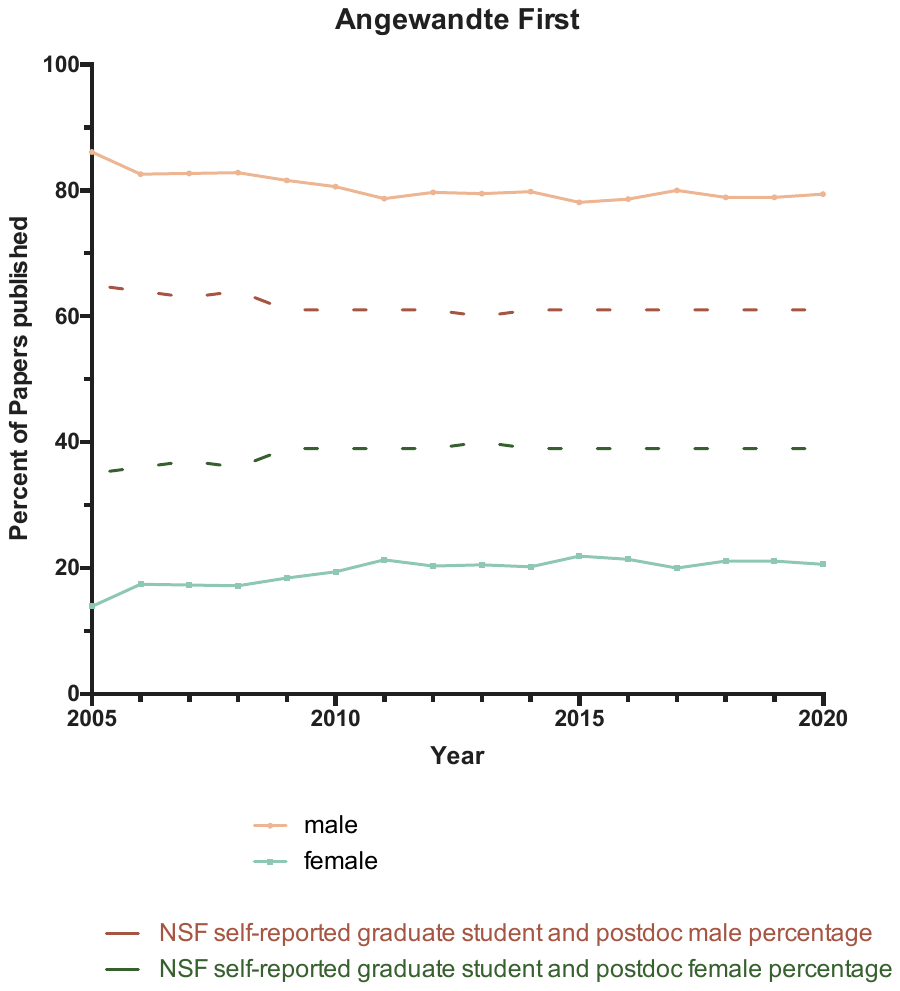

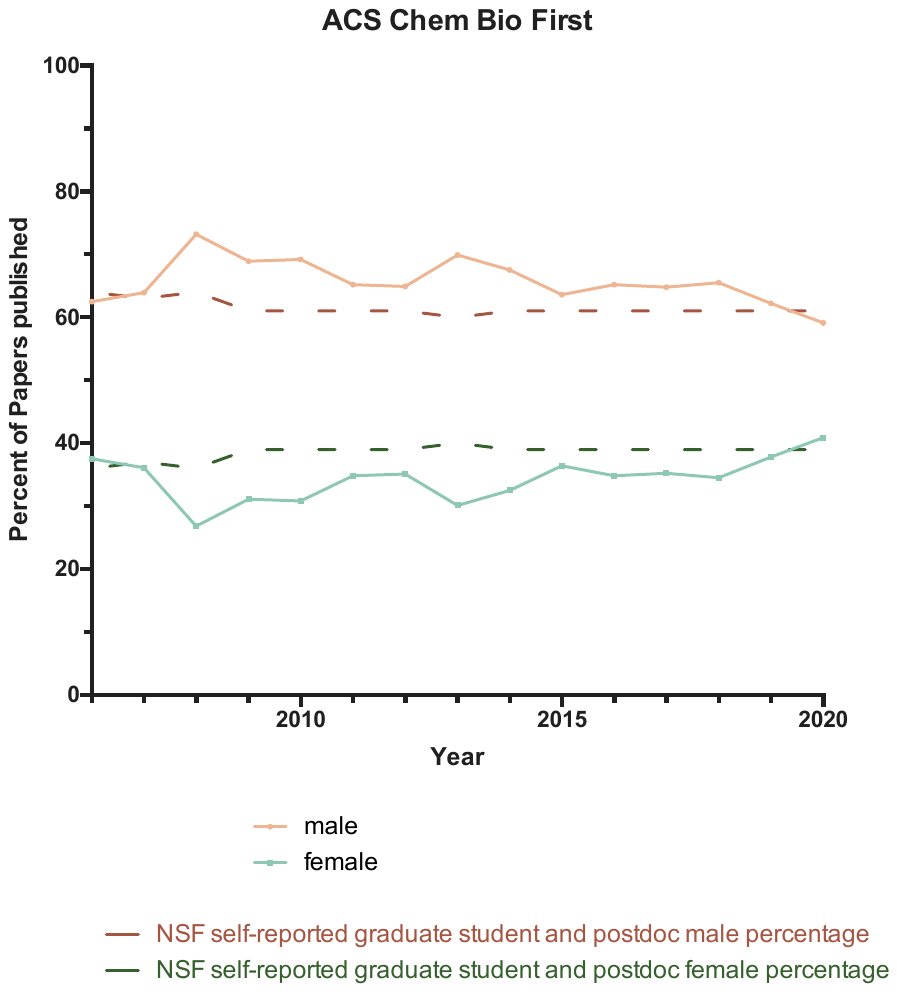

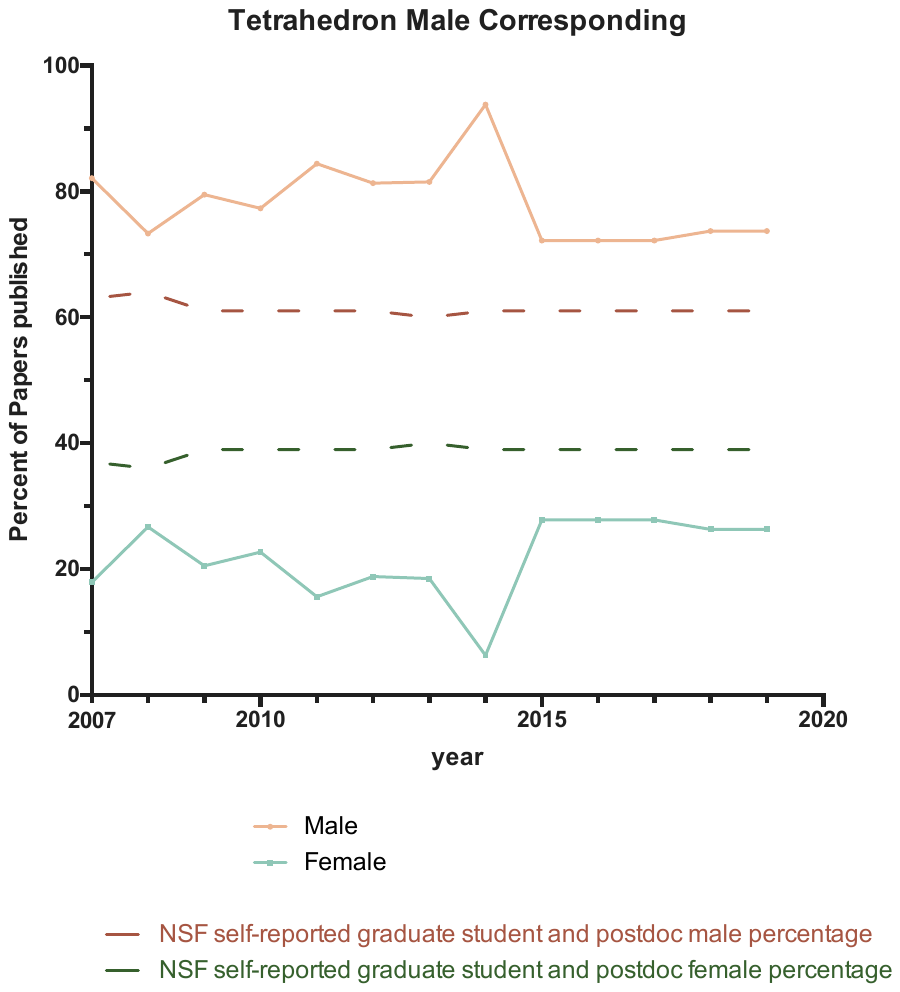
First author percentages for male corresponding authors by journal

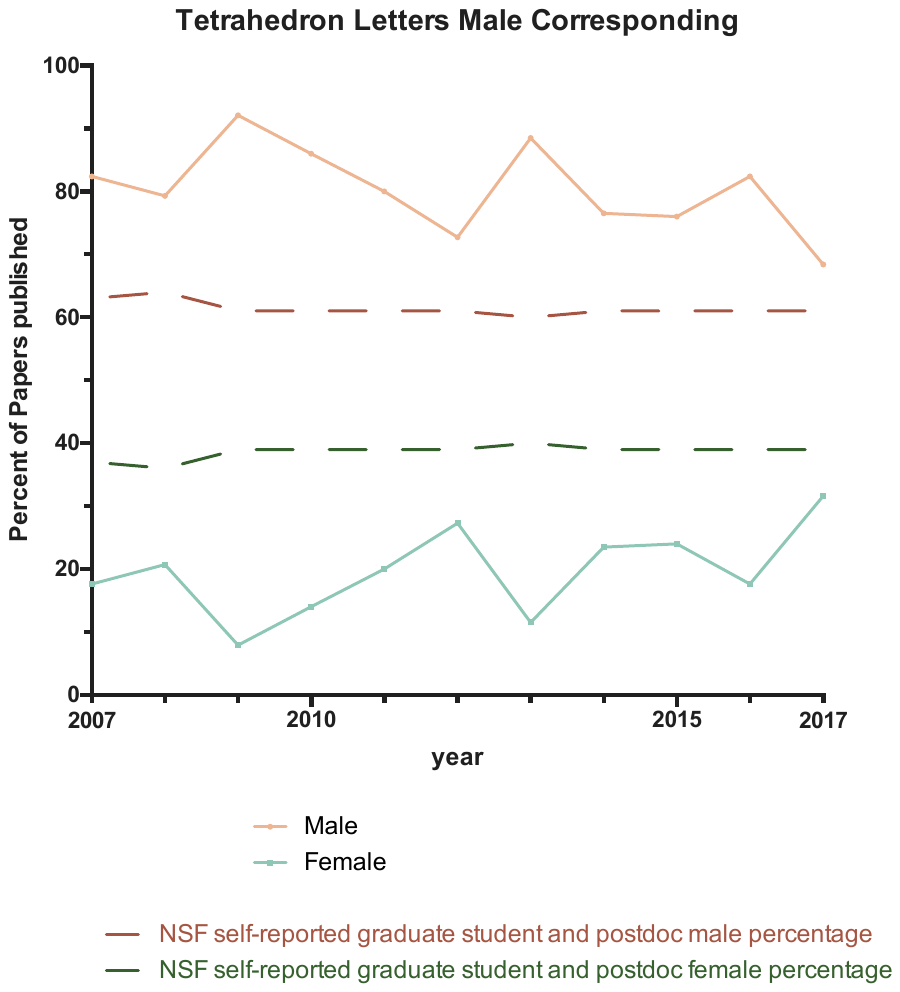

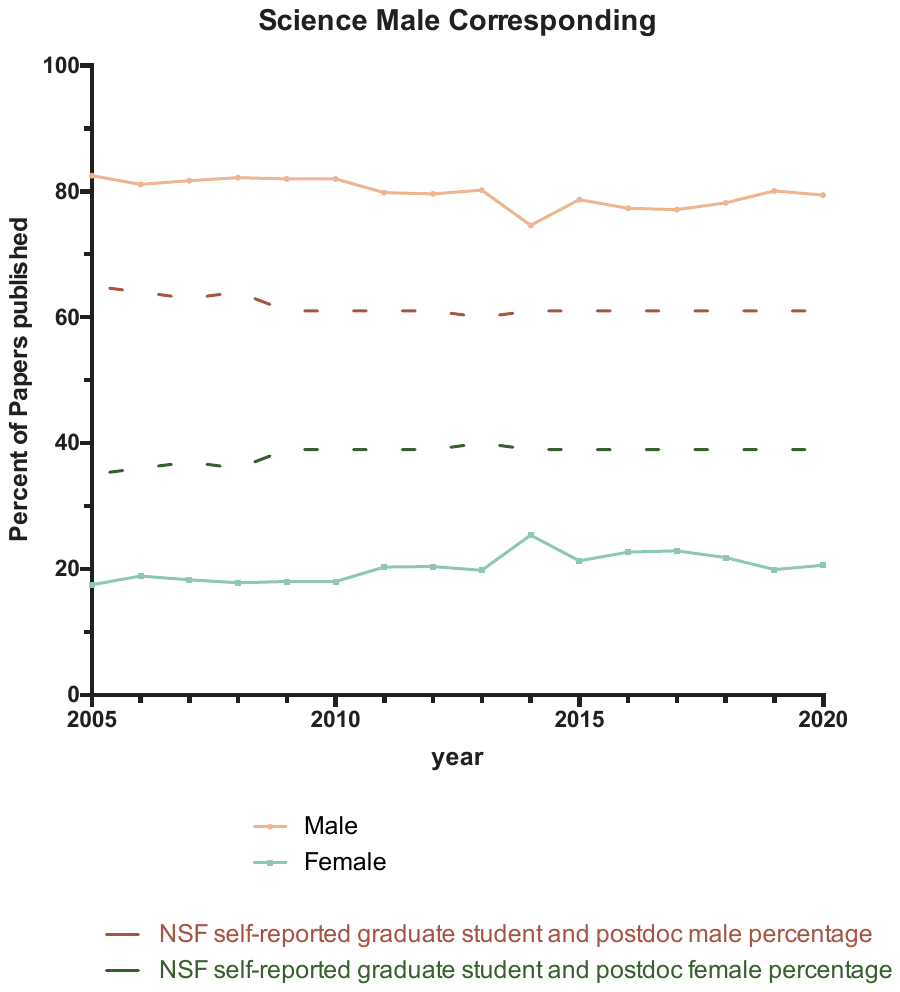

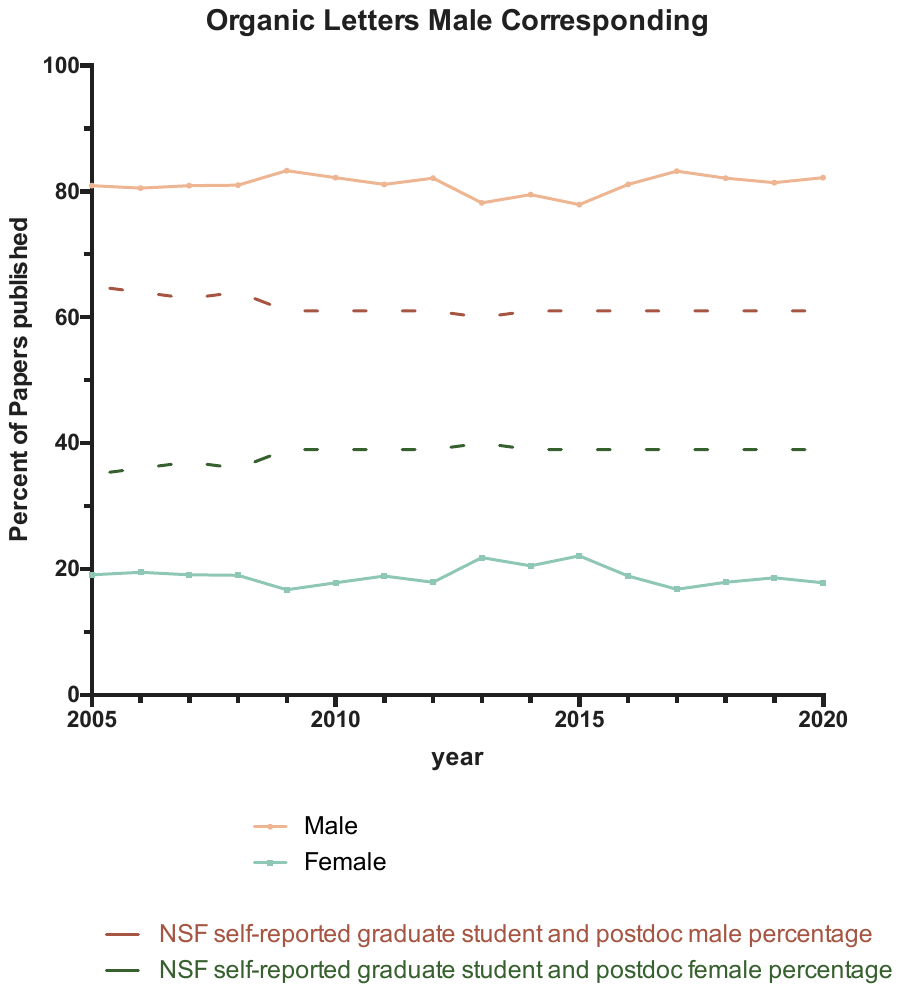

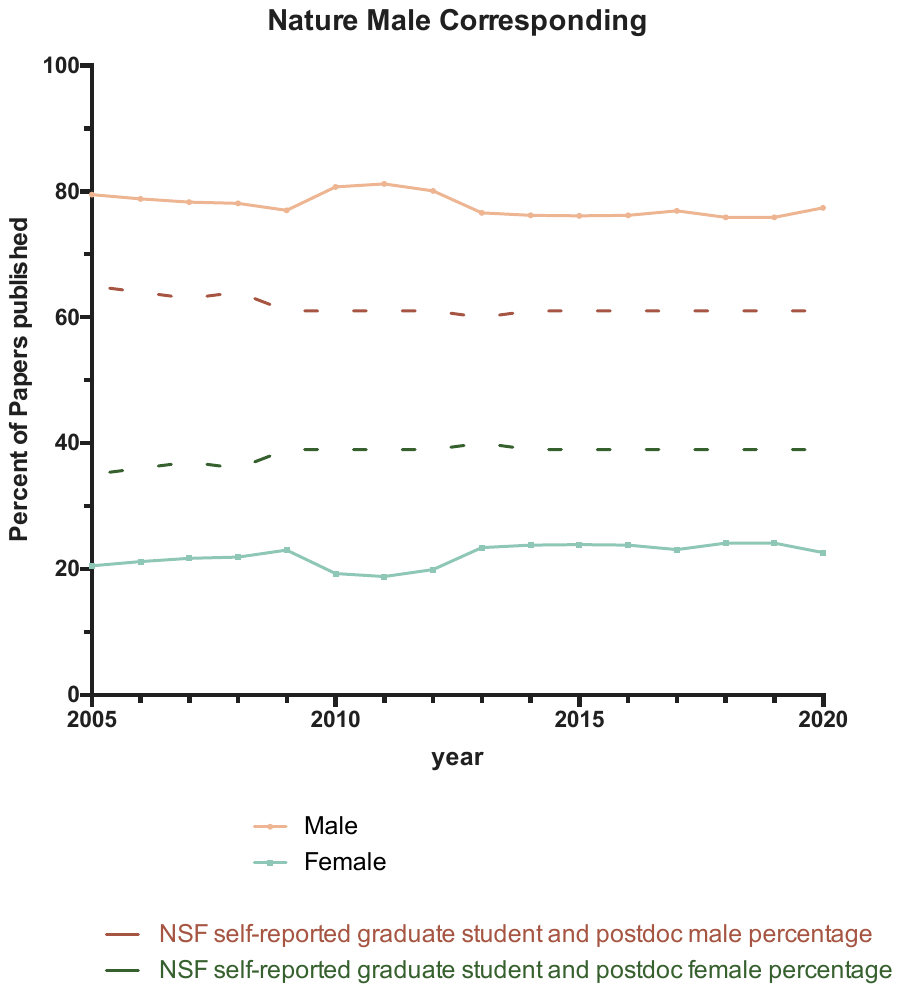

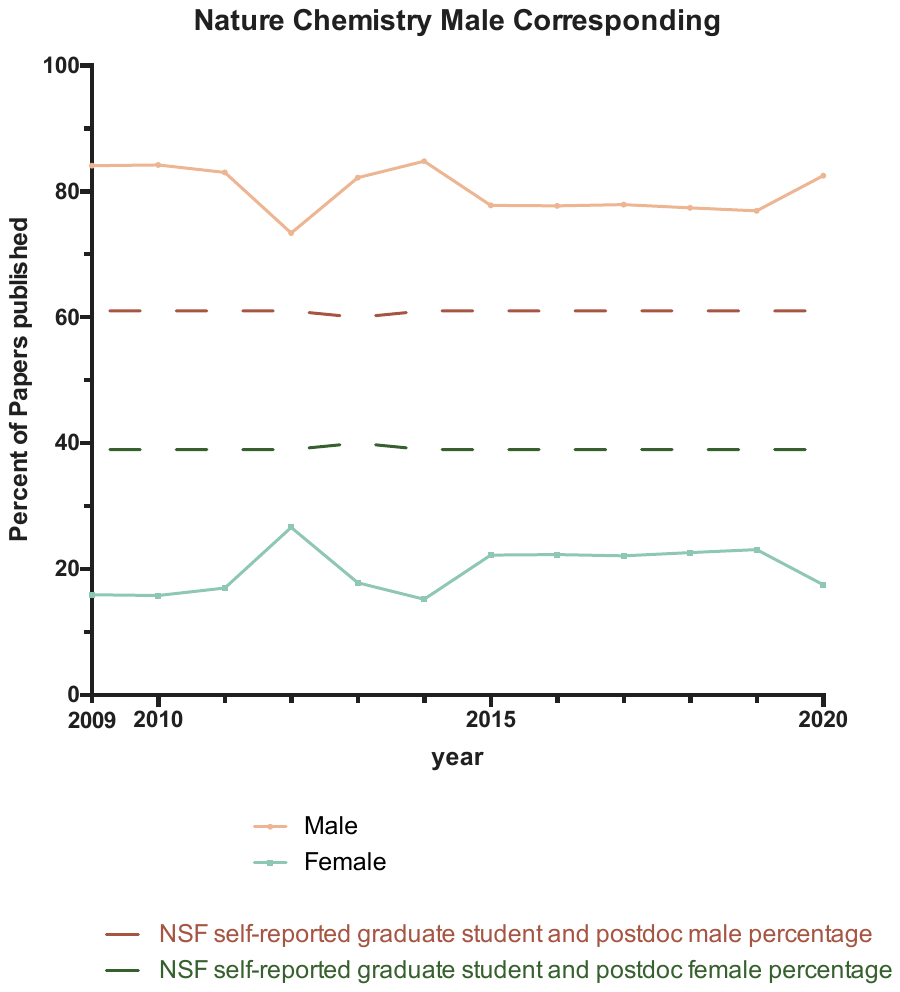

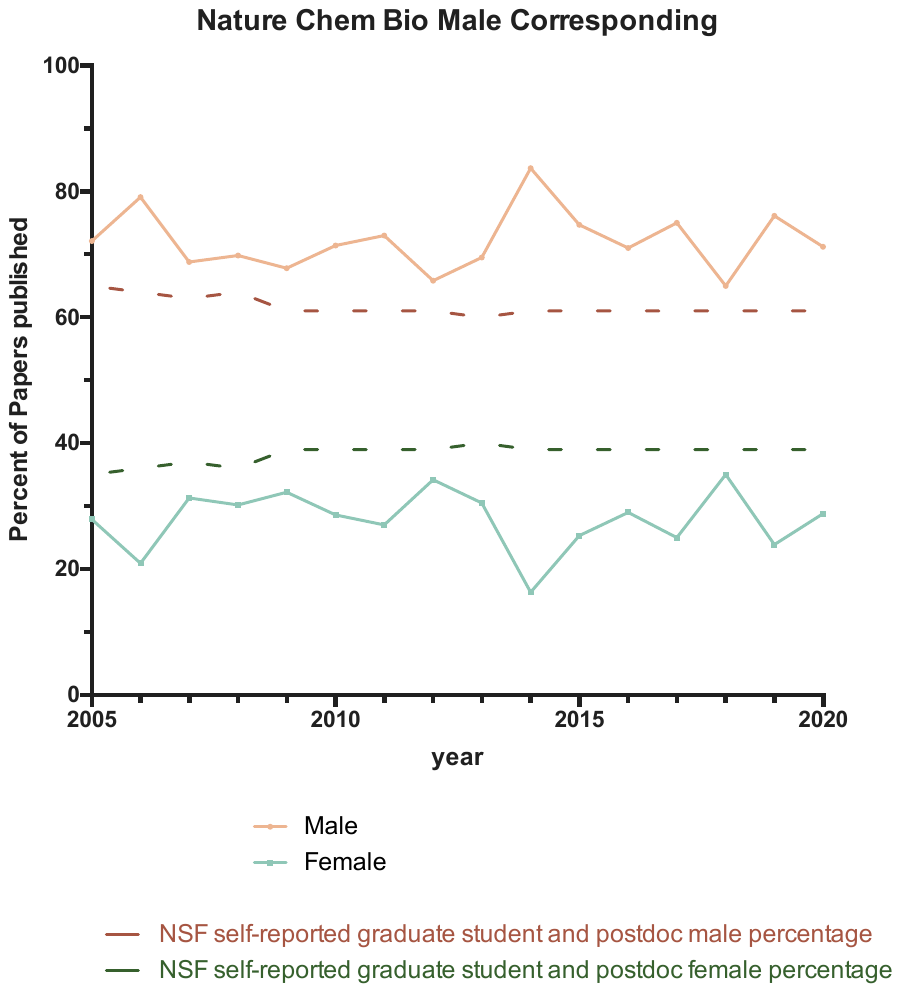

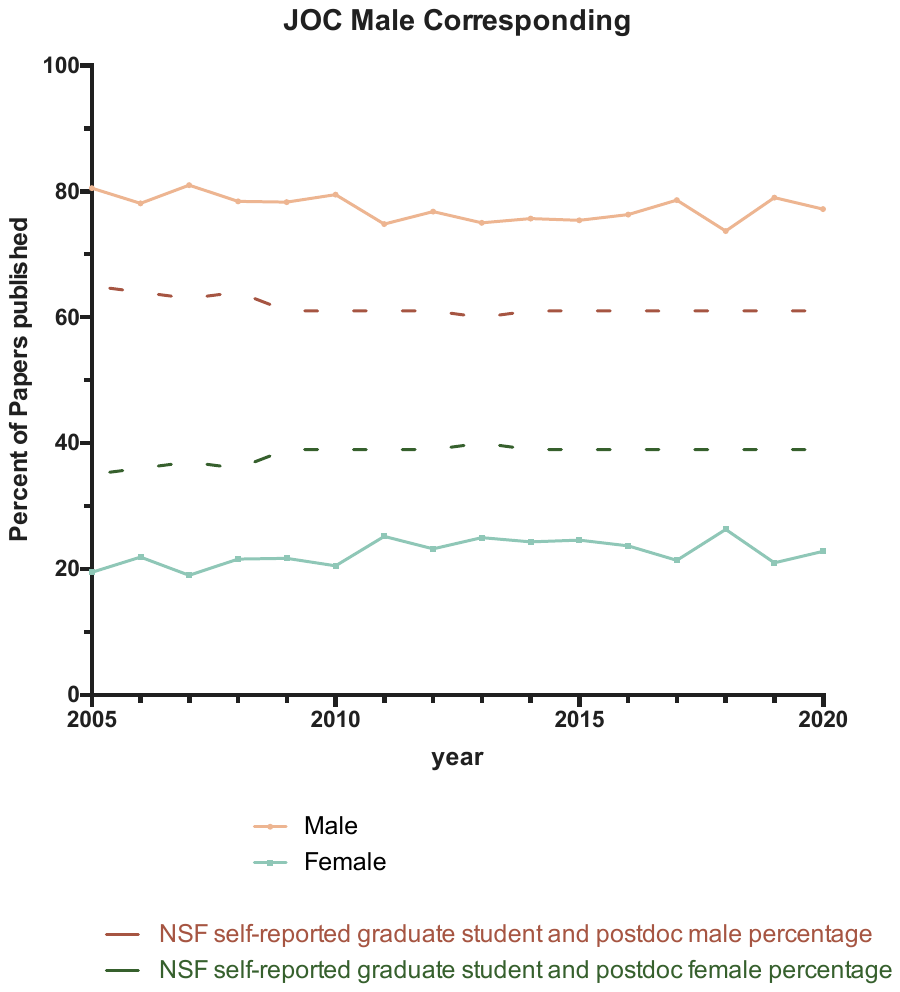

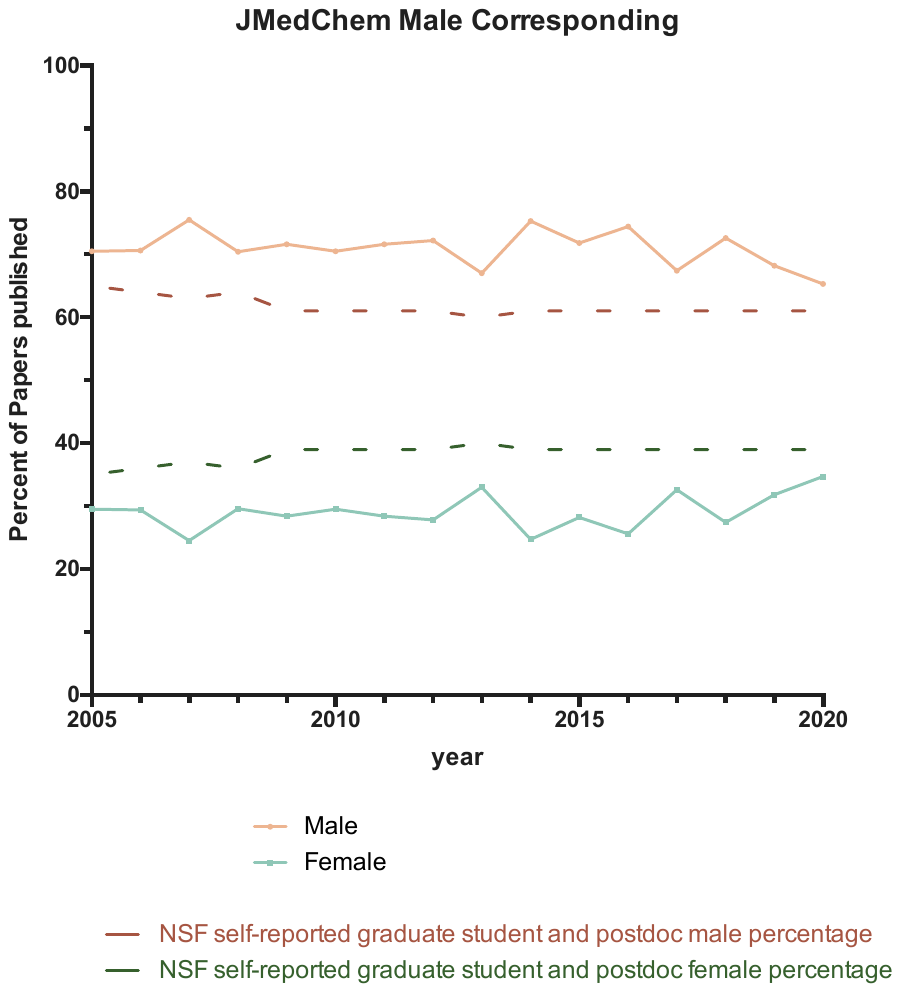

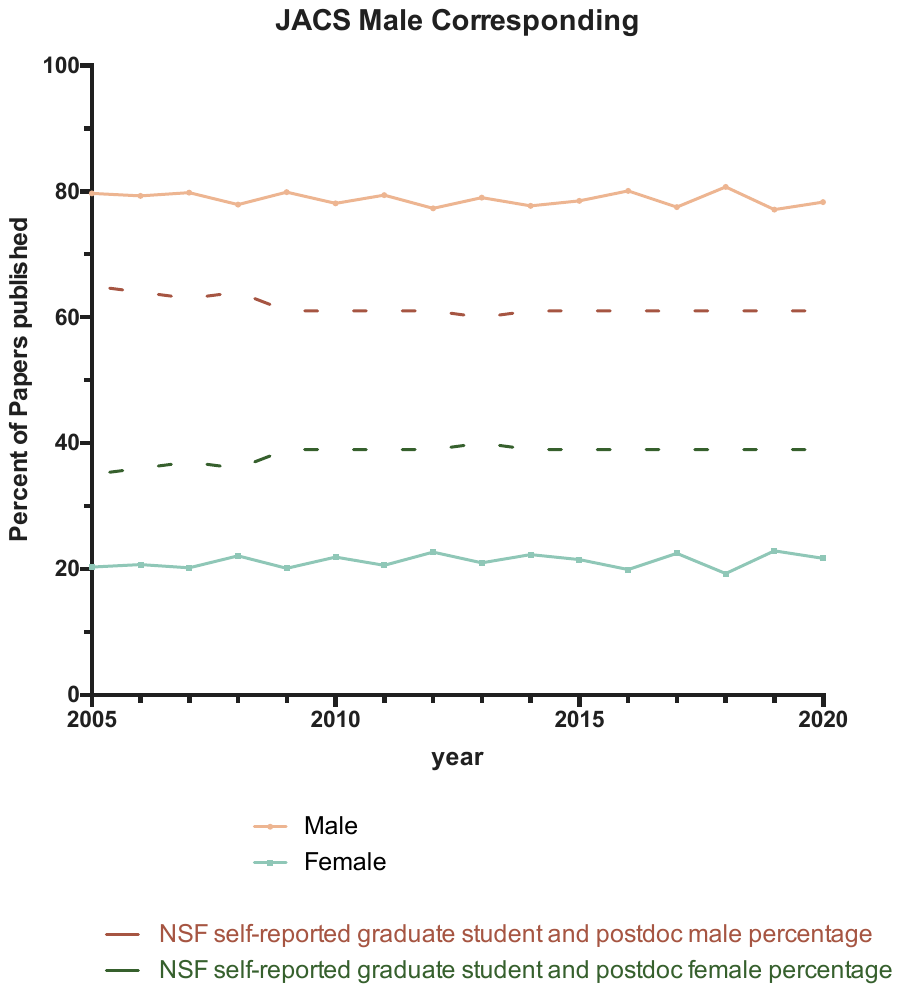

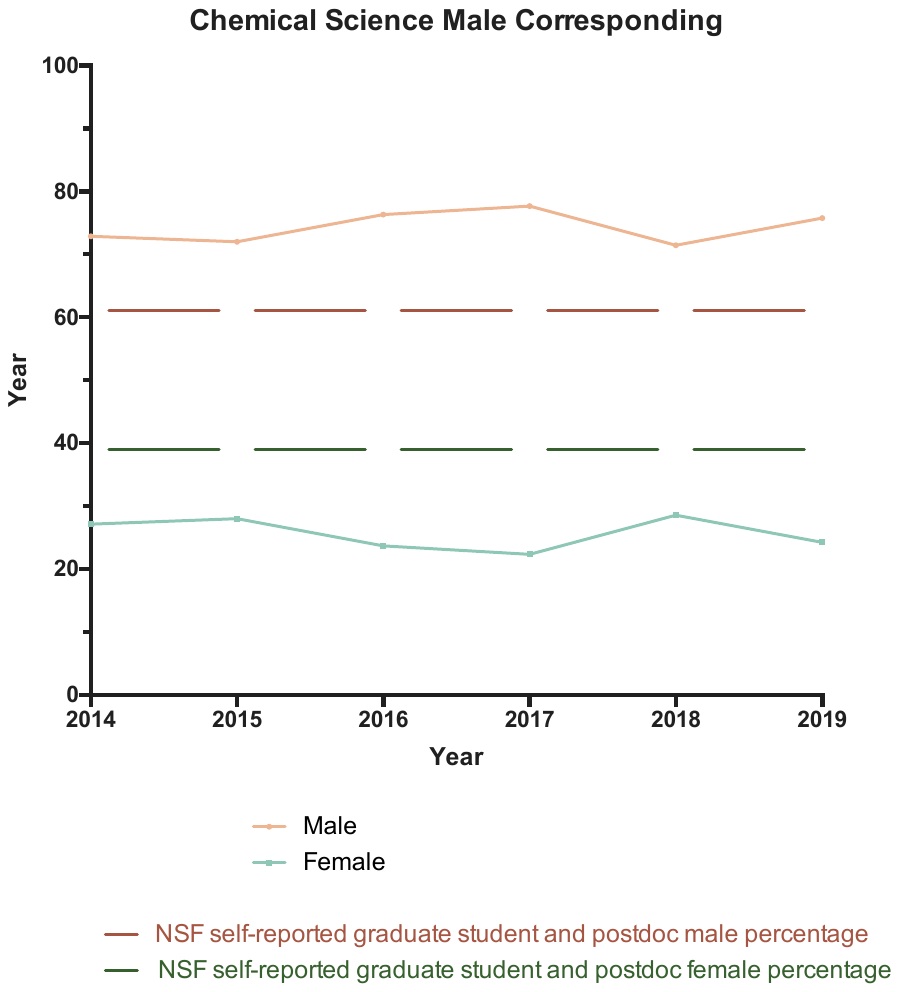

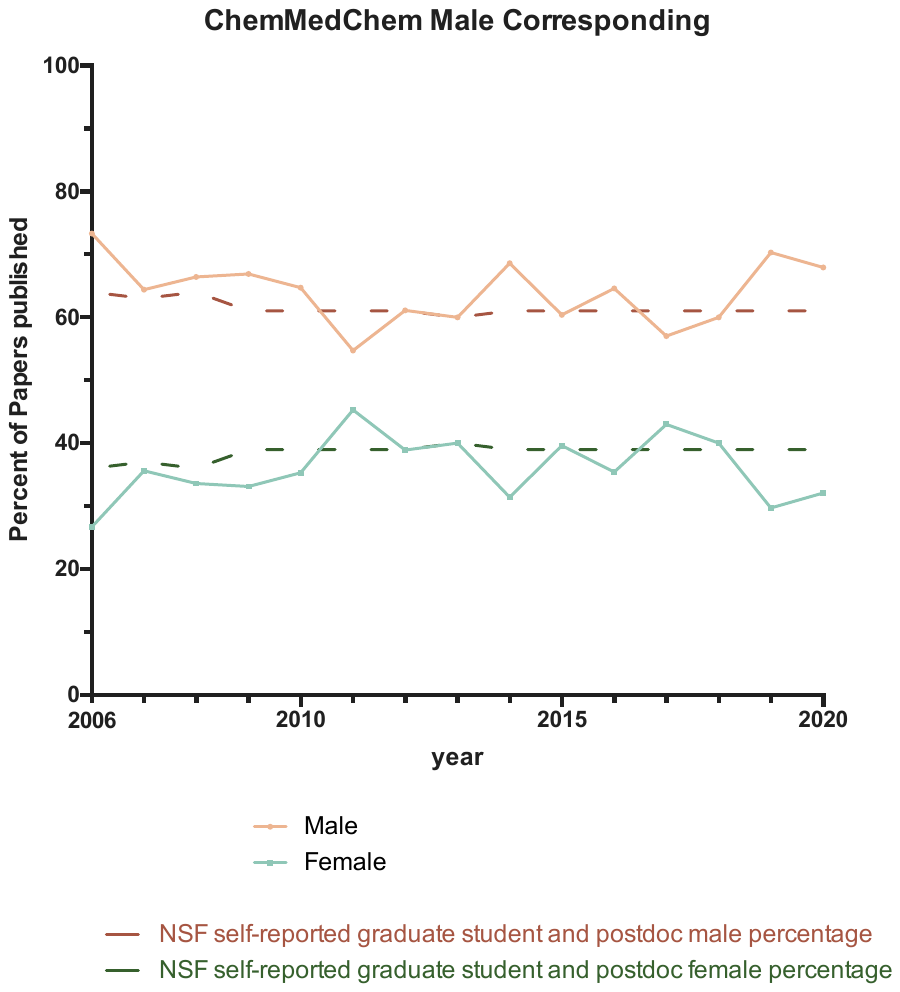

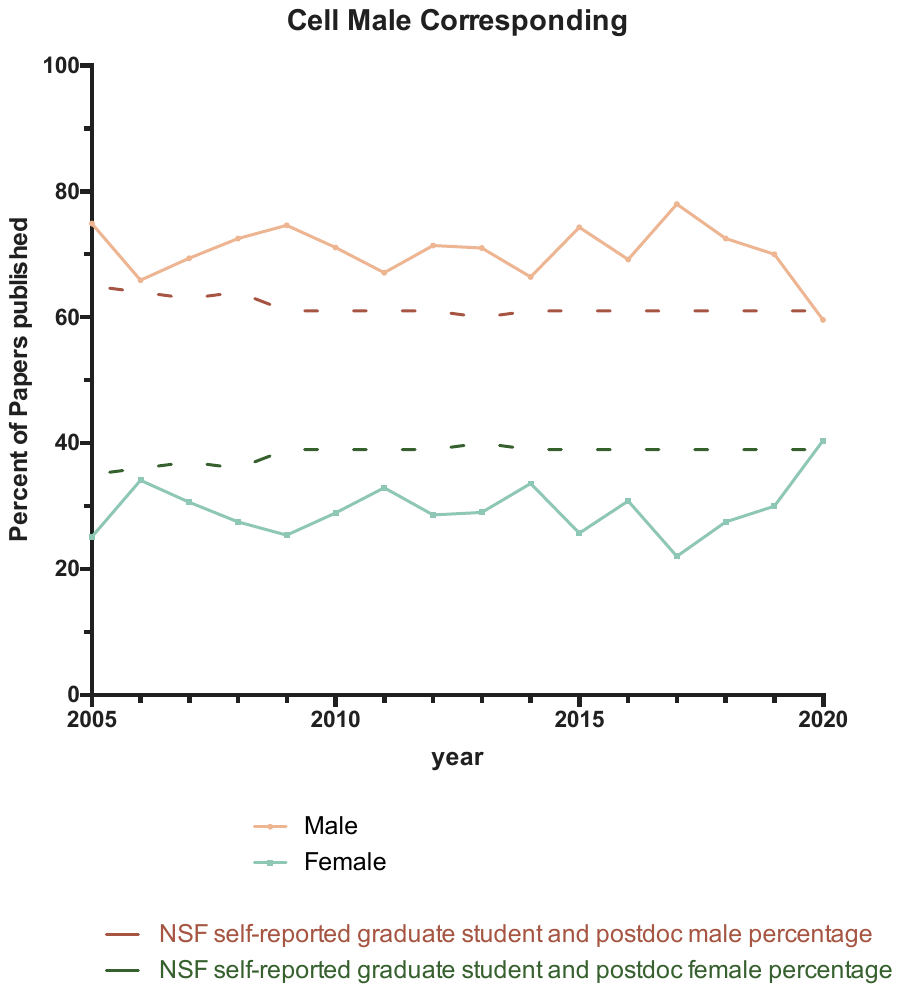

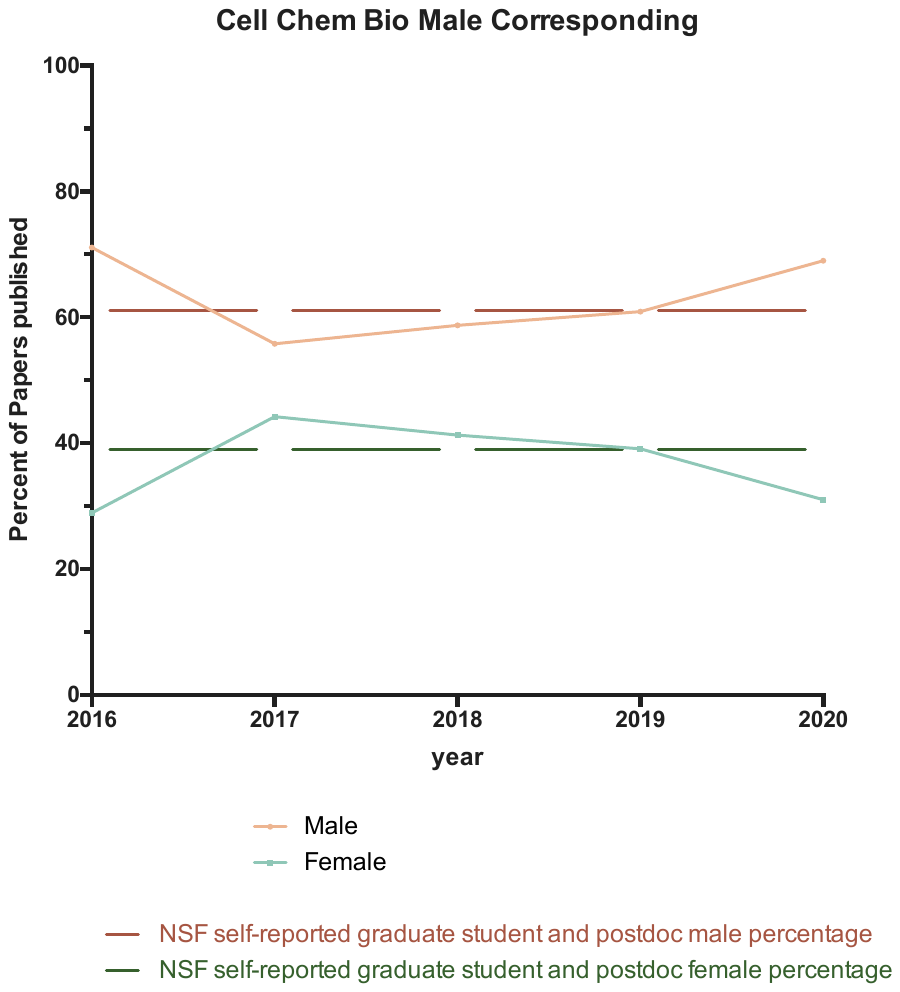

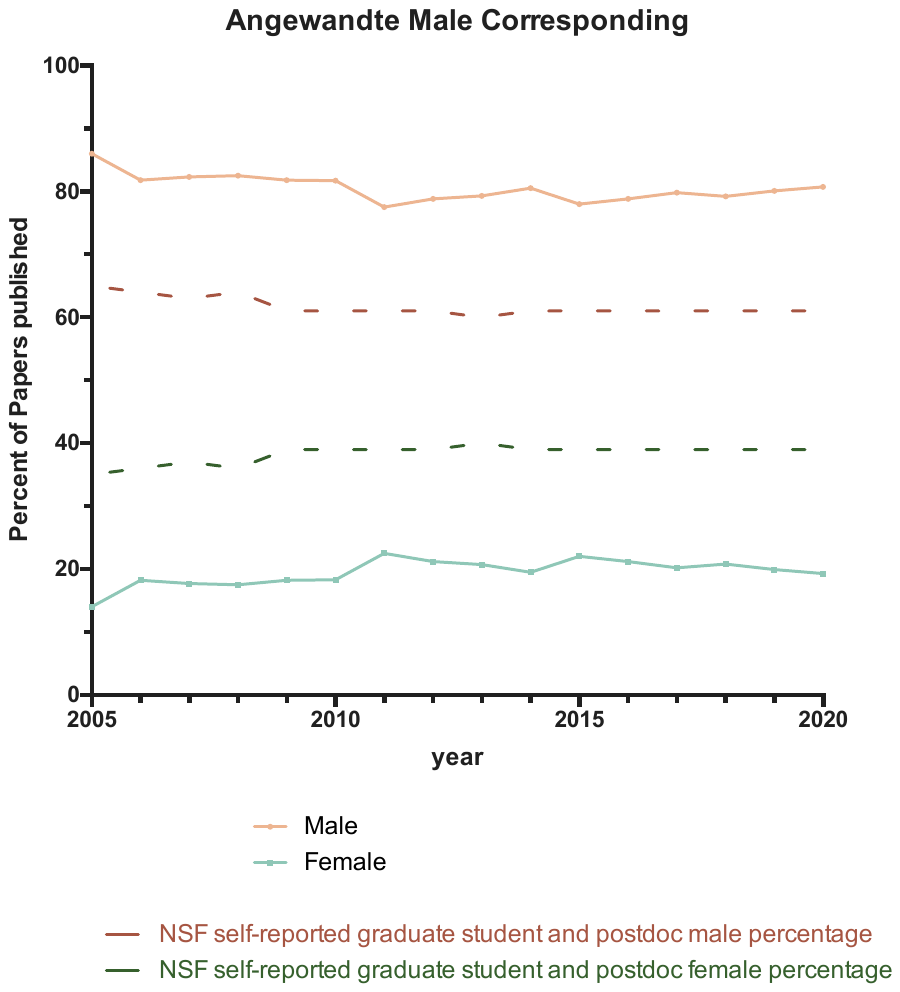

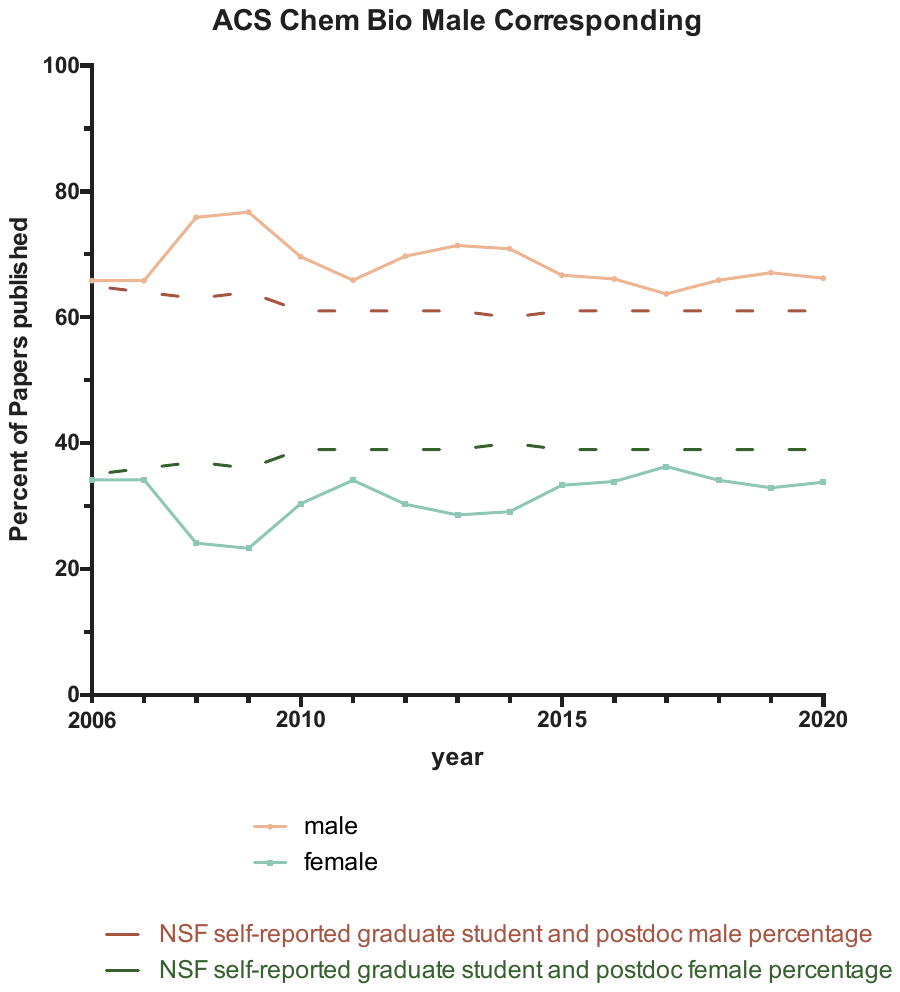

First author percentages for female corresponding authors by journal

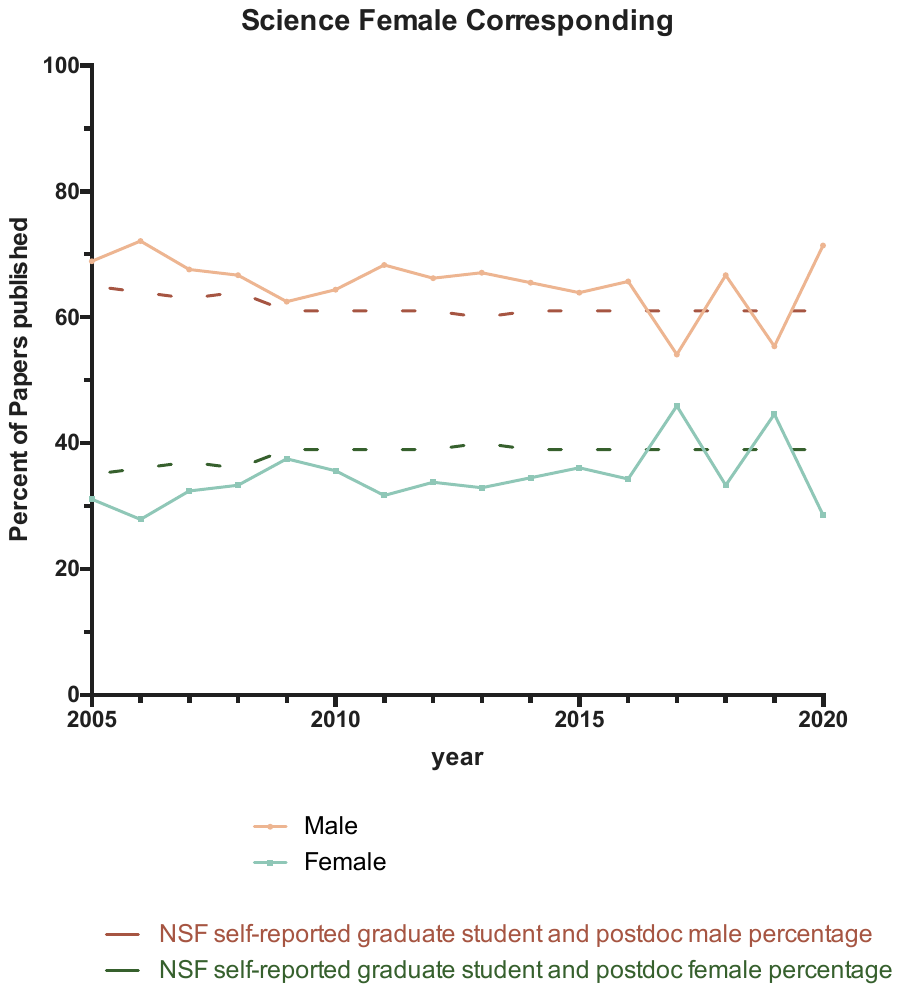

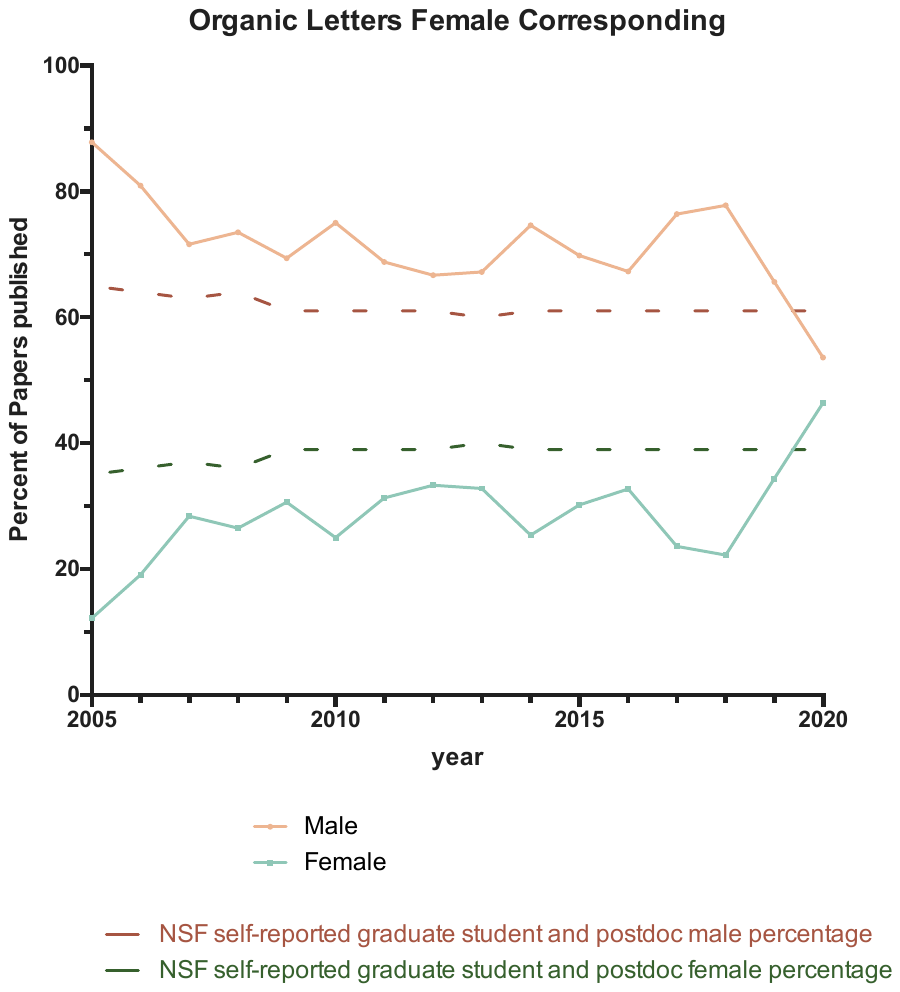

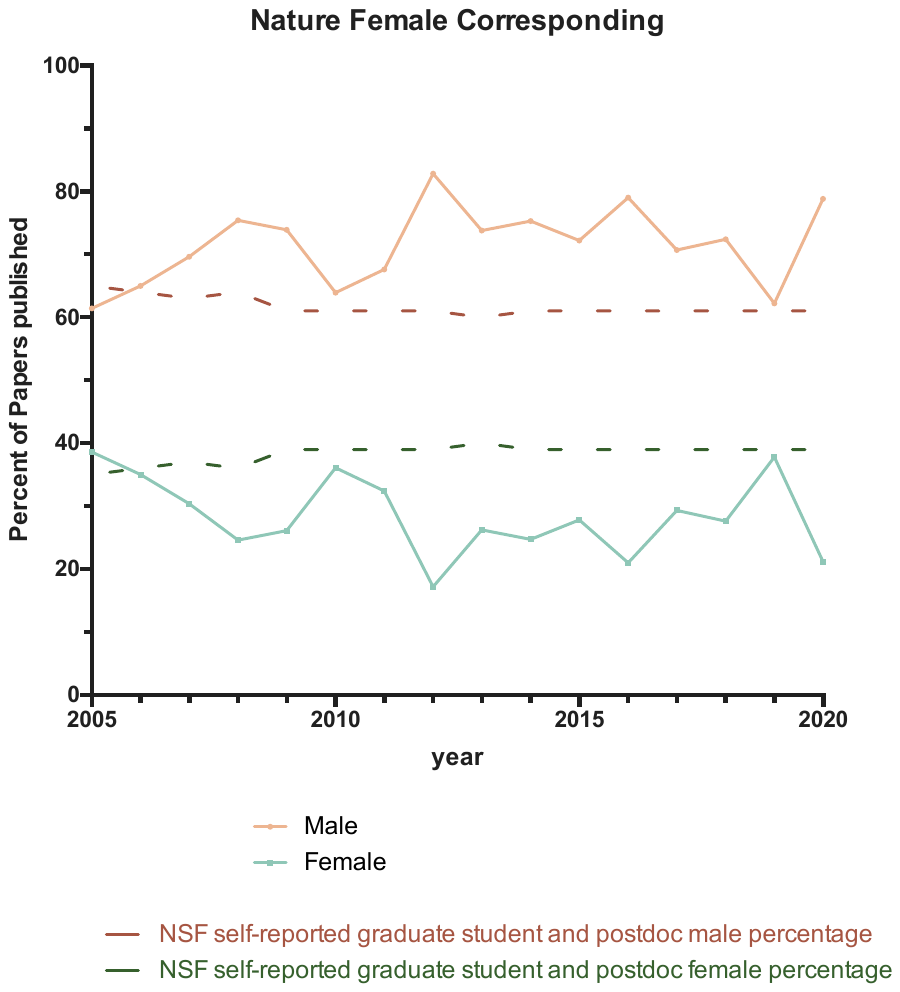

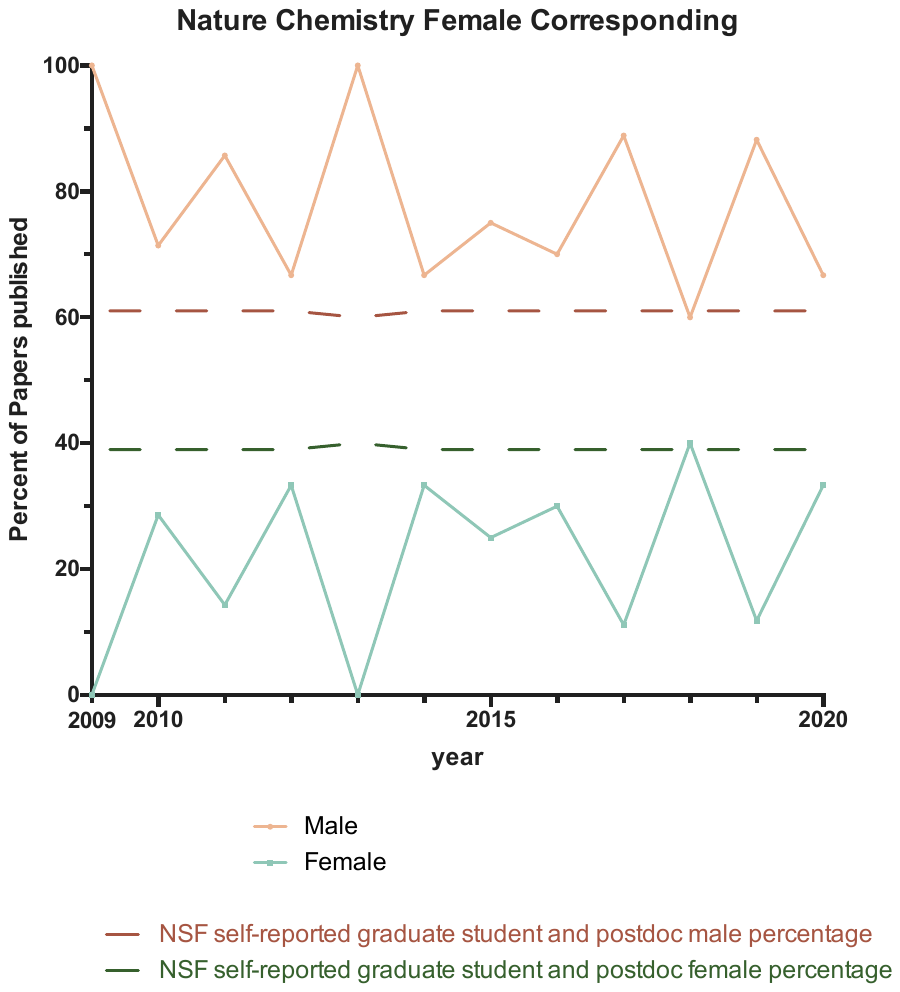

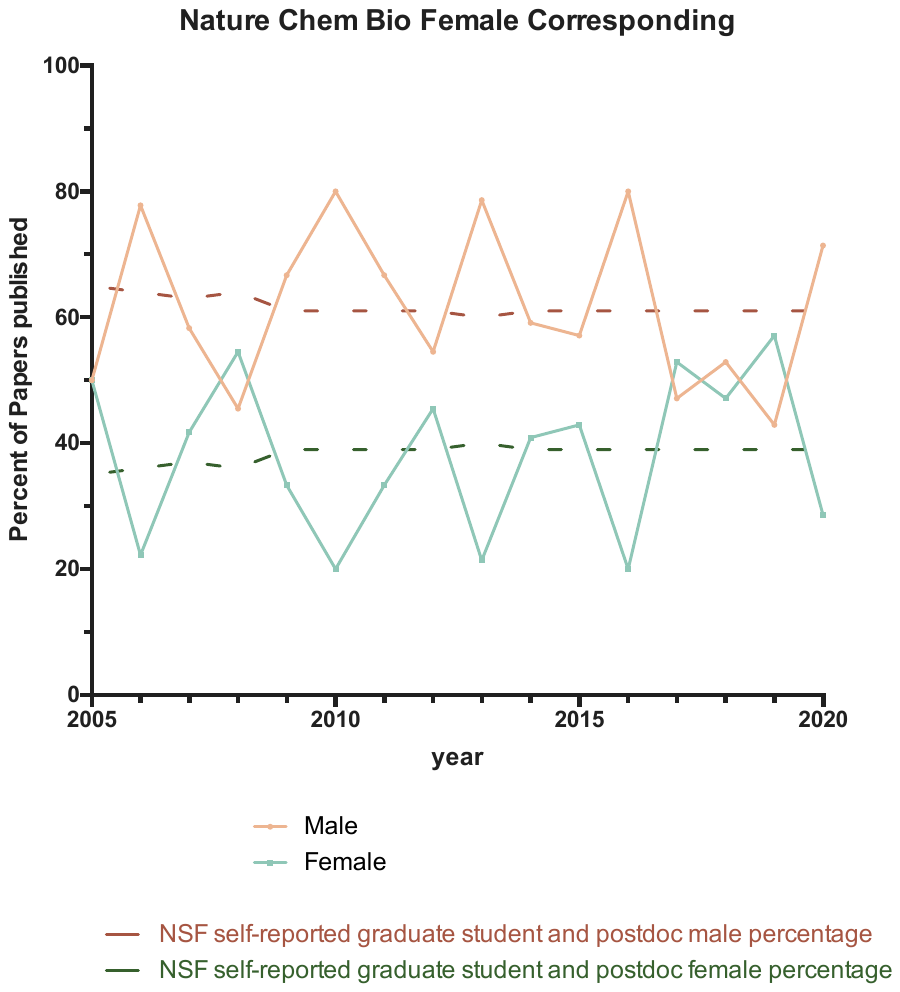

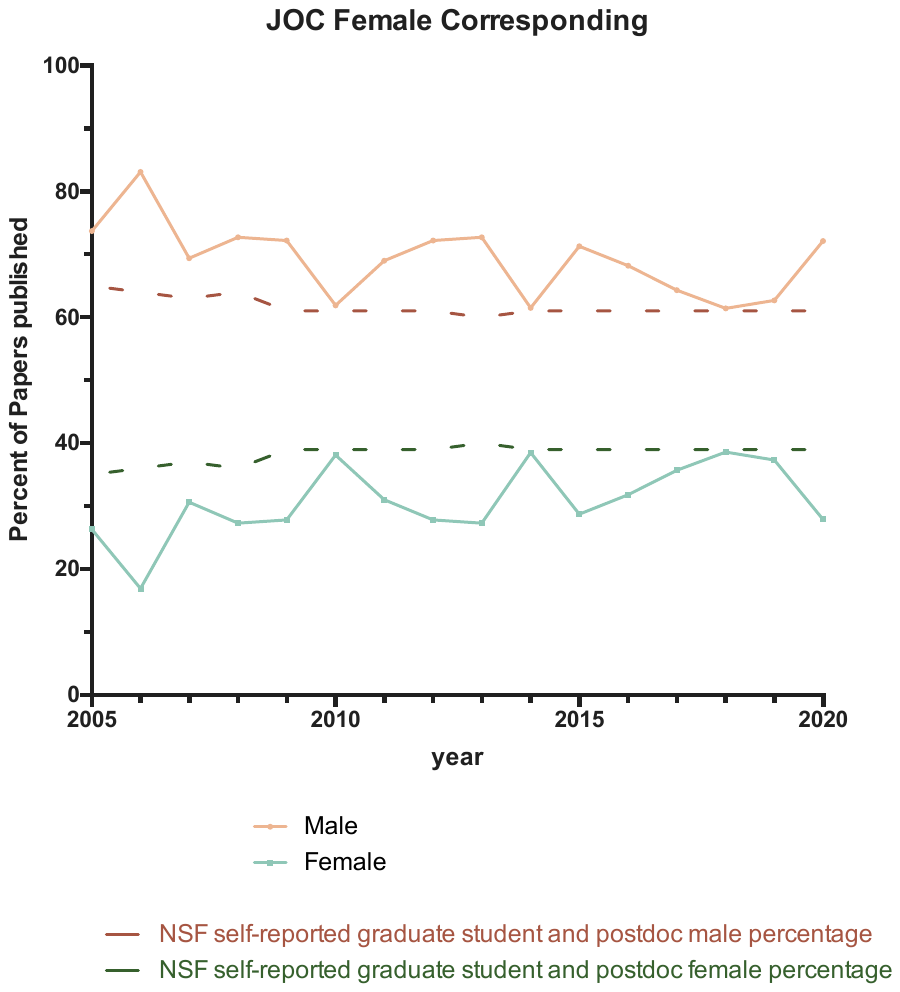

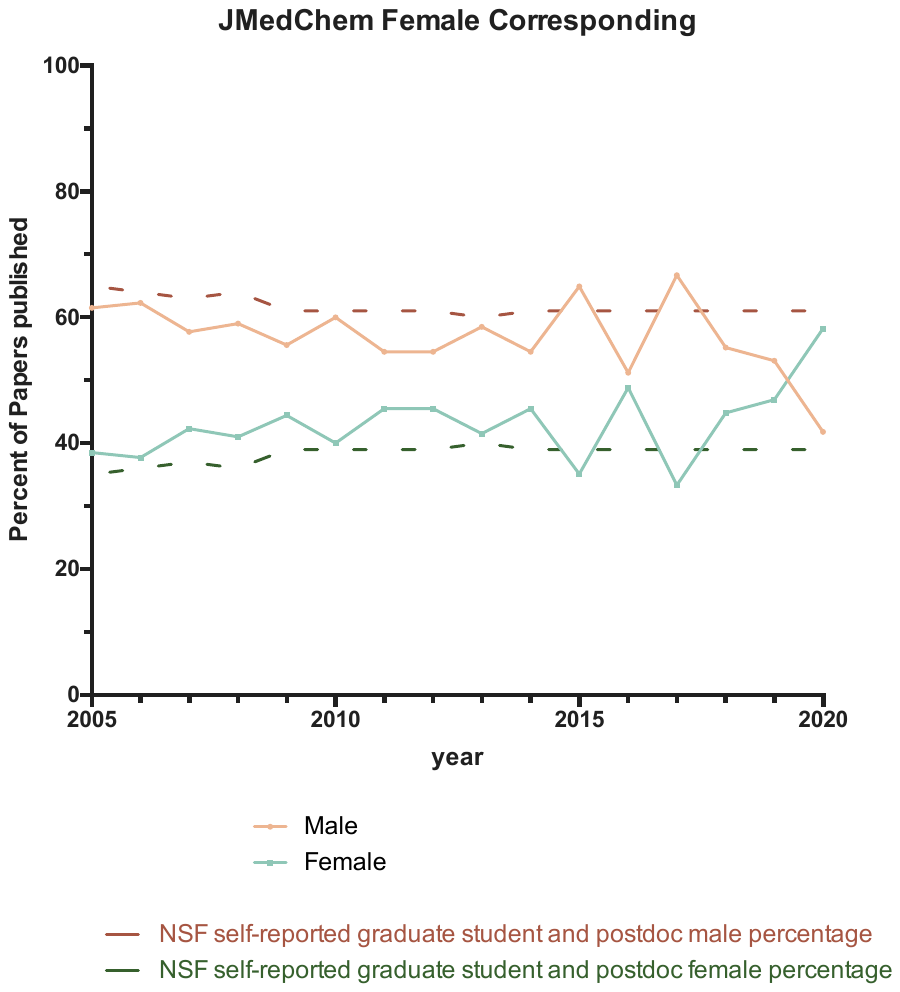

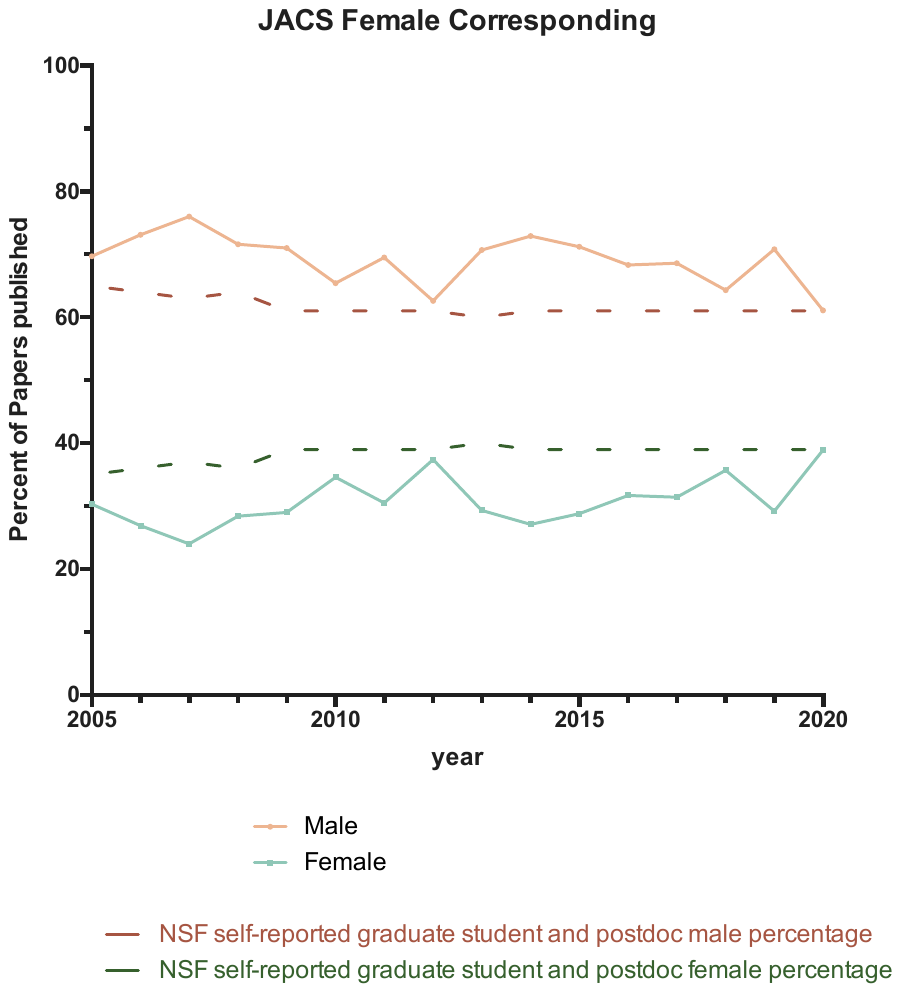

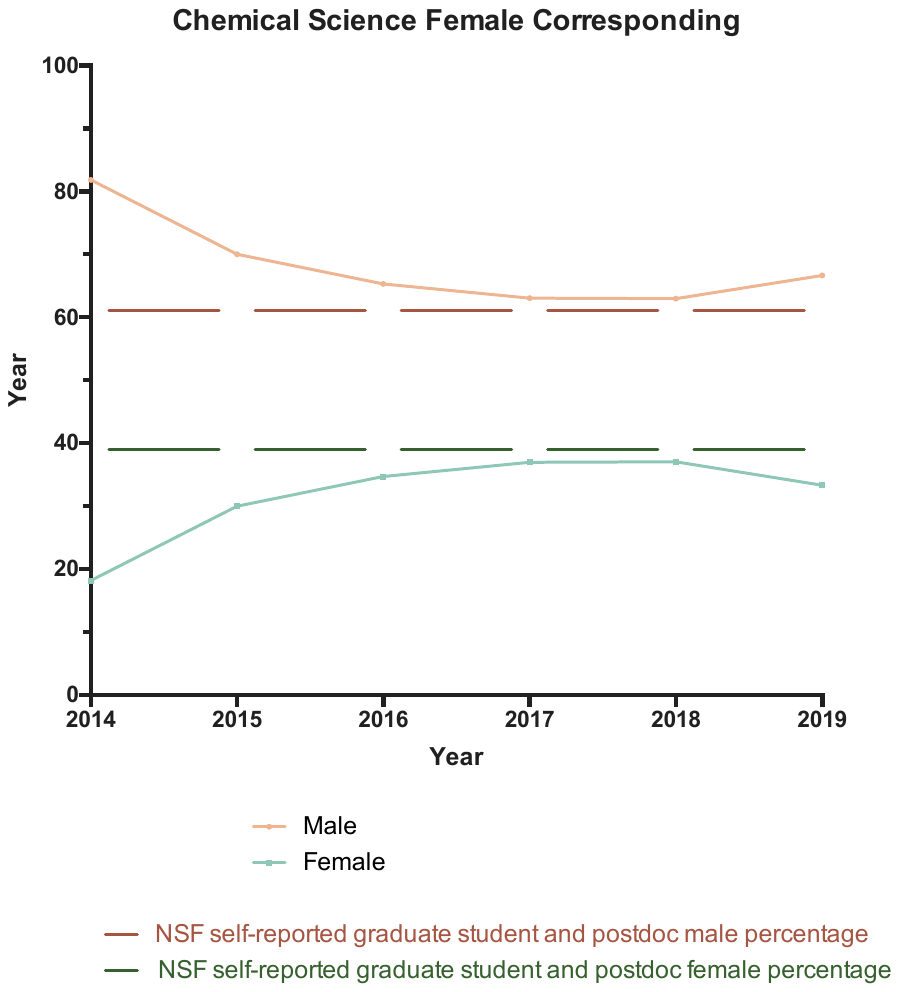

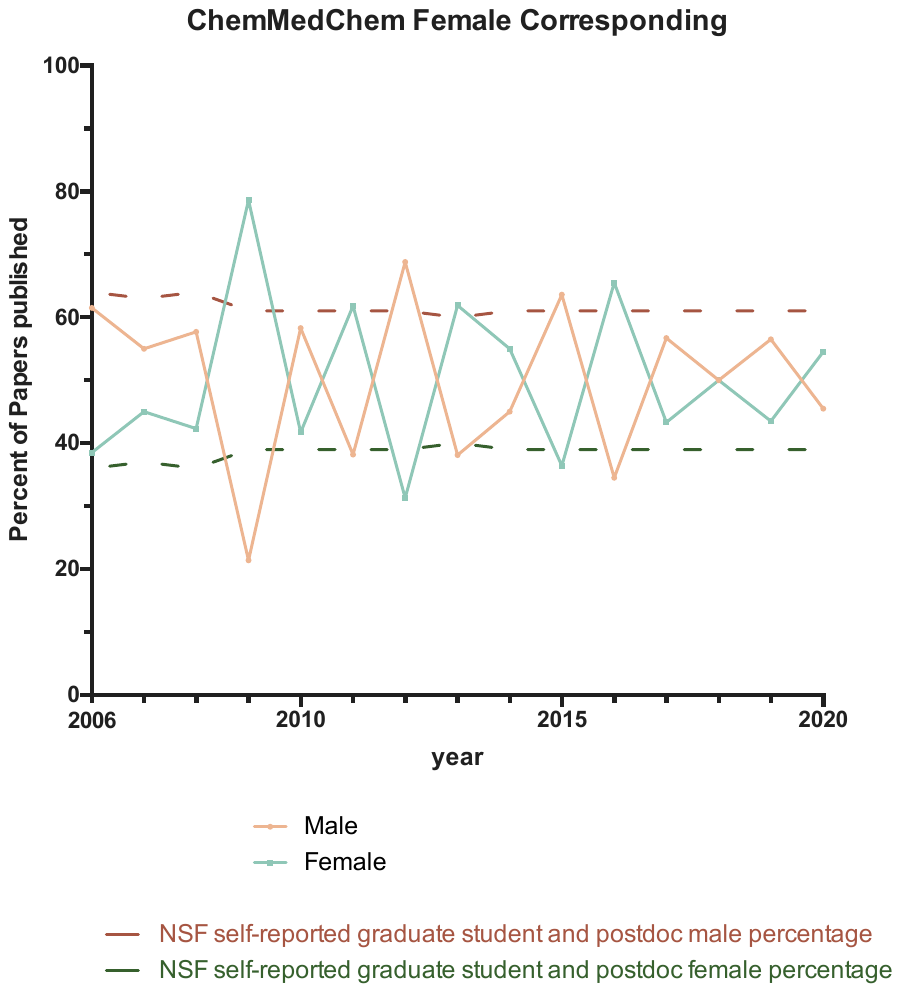

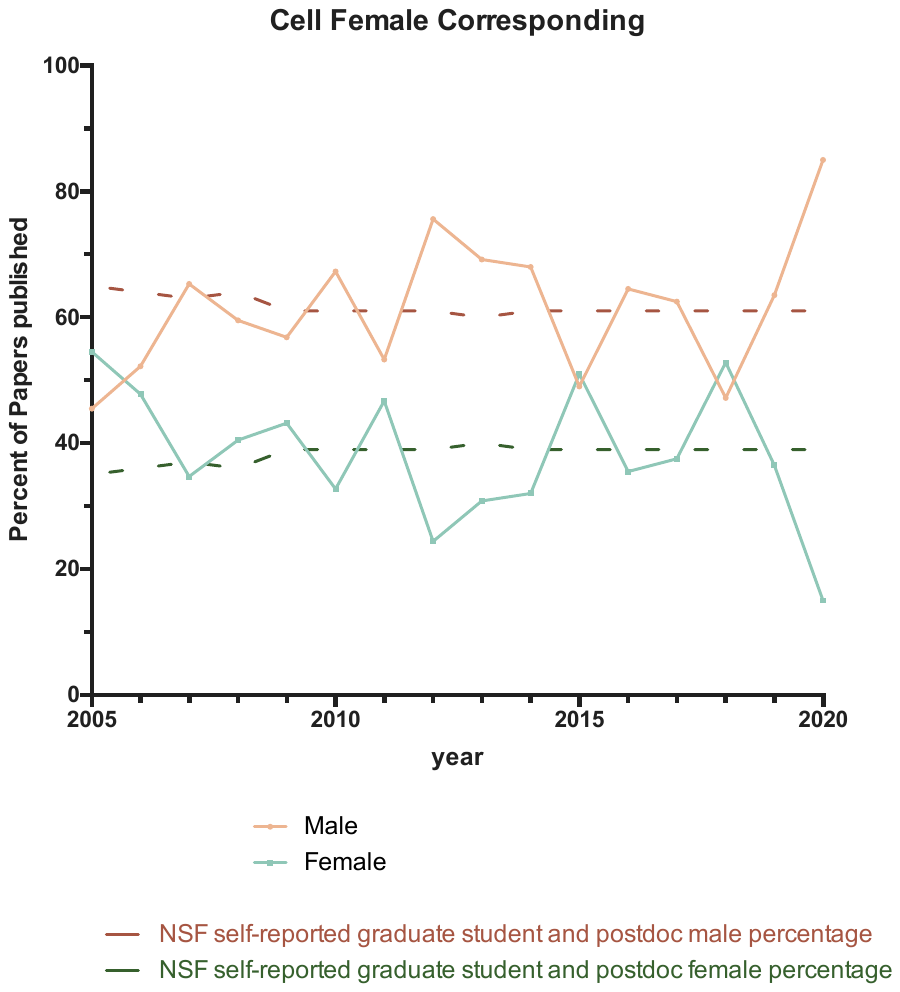

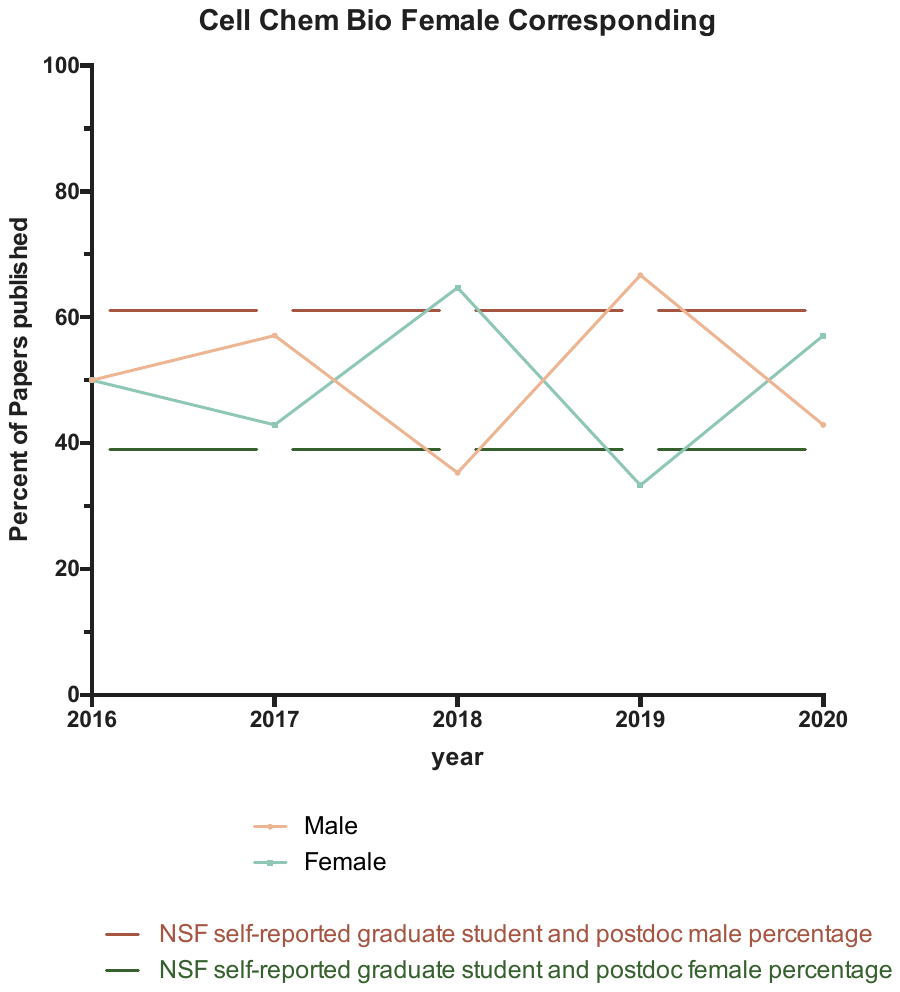

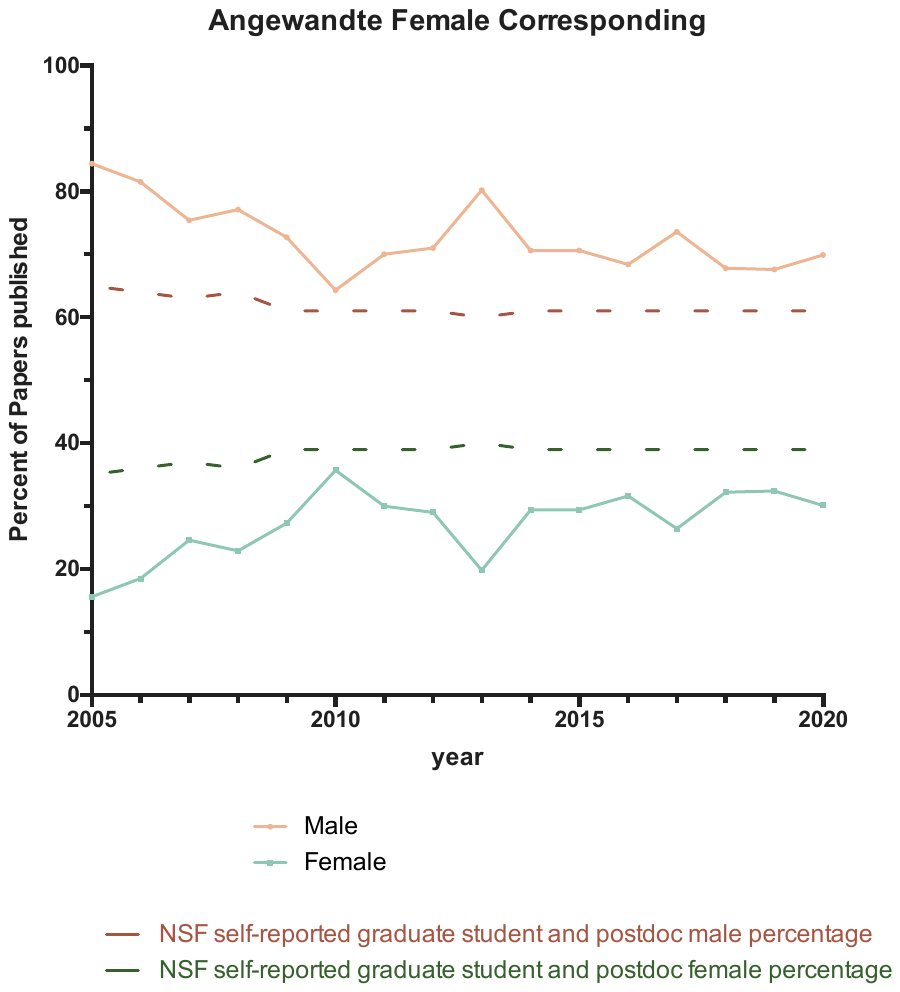

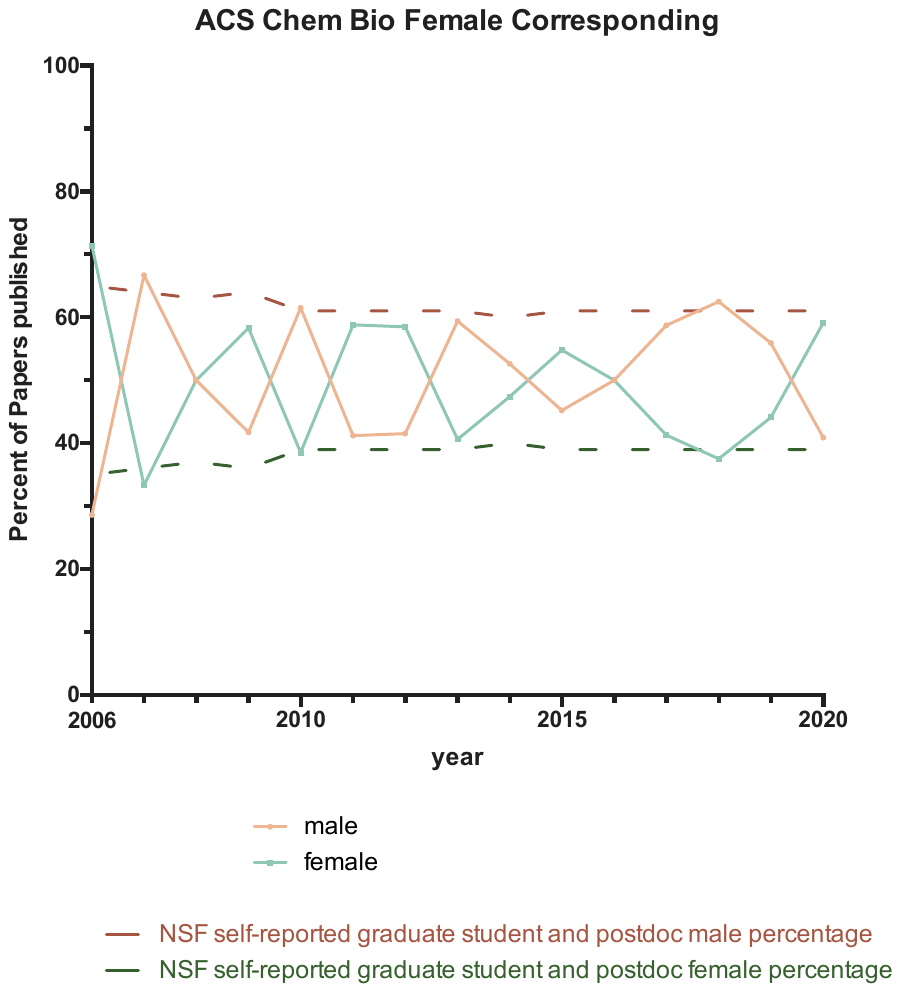
